# Predictability of cortico-cortical connections in the mammalian brain

**DOI:** 10.1101/2020.12.03.410803

**Authors:** Ferenc Molnár, Szabolcs Horvát, Ana R. Ribeiro Gomes, Mária Ercsey-Ravasz, Kenneth Knoblauch, Henry Kennedy, Zoltan Toroczkai

**Affiliations:** Department of Physics University of Notre Dame, Notre Dame, IN 46556, USA; Center for Systems Biology Dresden, Pfotenhauerstr. 108, 1307 Dresden, Germany; Max Planck Institute for Cell Biology and Genetics, Pfotenhauerstr. 108, 1307 Dresden, Germany; Max Planck Institute for the Physics of Complex Systems, Nöthnitzerstr. 38, 01187 Dresden, Germany; Univ Lyon, Université Claude Bernard Lyon 1, INSERM, Stem Cell and Brain Research Institute U1208, 69500 Bron, France; Section on Cognitive Neurophysiology and Imaging, Laboratory of Neuropsychology, National Institute of Mental Health, National Institutes of Health, Bethesda, MD, 20892, USA; Faculty of Physics, Babeş-Bolyai University, 400084 Cluj-Napoca, Romania; Transylvanian Institute of Neuroscience, 400157 Cluj-Napoca, Romania; Institute of Neuroscience, Center for Excellence in Brain Science and Intelligence Technology, Chinese Academy of Sciences, Shanghai 200031, China; Shanghai Center for Brain Science and Brain-Inspired Intelligence Technology, Shanghai 200031, China

**Keywords:** machine learning, neuroanatomy, neocortex, primate, rodent

## Abstract

Despite a five-order magnitude range in size, the mammalian brain exhibits many shared anatomical and functional characteristics that should translate into cortical network commonalities. Here we develop a framework employing machine learning to quantify the degree of predictability of the weighted interareal cortical matrix. Data were obtained with retrograde tract-tracing experiments supplemented by projection length measurements. Using this framework with consistent and edge-complete empirical datasets in the macaque and mouse cortex, we show that there is significant amount of predictability embedded in the interareal cortical networks of both species. At the binary level, links are predictable with an Area Under the ROC curve of at least 0.8 for the macaque. At the weighted level, strengths of the medium and strong links are predictable with at least 85-90% accuracy in mouse and 70-80% in macaque, whereas weak links are not predictable in either species. These observations suggest that the formation and evolution of the cortical network at the mesoscale is to a large extent, rule-based, motivating further research on the architectural invariants of the cortical connectome.

## Introduction

Information in the brain is encoded via the temporal patterns of signals generated by a network of distributed neuronal assemblies (Hebb, 1949; McCulloch and Pitts, 1943), whose organization has been shown to be strongly determined by its weighted connectivity and spatial embedding (Knoblauch et al., 2016; Markov et al., 2013a). This contrasts with technological information networks, where information including the destination address is encoded into packets and routed via switches, with the network structure serving merely as propagation backbone. In comparison, the structure of brain networks is considerably more complex and forms an integral part of its processing algorithm, the deciphering of which crucially hinges on the details of its connectome (Sporns et al., 2005). This is supported, for example, by the observation that many neurodegenerative diseases stem from neuronal pathway disruptions (Delbeuck et al., 2007; Friston and Frith, 1995; Peters et al., 2013; Silva et al., 2015).

Despite being fundamental for understanding the brain in health and disease, there is limited knowledge of cortical circuitry, which at the microscale is presently intractable, due to the staggering size of its numbers of nodes (neurons) and connections (Frégnac and Bathellier, 2015). What is tractable with current technology, however, is the investigation of the meso-scale, interareal network corresponding to the pathways between functionally defined areas, addressed in ongoing electrophysiology and whole brain imaging efforts to understand cognitive functions (Mesulam, 2012). While the full interareal network (FIN) is currently unavailable for any mammal, it is obtainable in the foreseeable future, although, nevertheless, requiring highly specialized laboratories.

Among the empirical approaches, retrograde tract-tracing has emerged as a reliable, high-resolution method to trace neuronal pathways (Köbbert et al., 2000; Lanciego and Wouterlood, 2011). Compared to anterograde techniques, the major advantage of retrograde tract-tracing is that counts of labeled cells provide a reliable metric of connection strength, yielding a weighted, directed and spatially embedded, physical network of connections between brain areas (Gămănuţ et al., 2018; Majka et al., 2020; Markov et al., 2014; Zingg et al., 2014). In these experiments a single area (referred to as the target area) is injected with a tracer, which then back-labels the cell-bodies of neurons with terminals ending in the target area. Areas external to the target area housing labeled neurons are referred to as source areas. The weight of an interareal connection from area *j* to area *i*, defined via the counts of labeled neurons, is recorded as the Fraction of Labeled Neurons *FLN_ij_* found in area *j (j ≠ i)*, when injecting into area *i*.

Existing retrograde tracing datasets do not have full network connectivity information; they do provide edge-complete subgraphs, i.e., networks formed by a subset of vertices whose connectivity within this subset is fully known. These studies show that interareal cortical networks (Majka et al., 2020) are not random graphs, but complex networks with characteristic structural features (Ercsey-Ravasz et al., 2013; Gămănuţ et al., 2018; Horvát et al., 2016; Theodoni et al., 2020). Moreover, interareal networks appear to be in a class of their own when compared to other real-world complex networks, including technological information networks (Milo et al., 2004). One of the most distinguishing feature of interareal networks is their high density of binary connectivity (connections existing or not), i.e., containing a large fraction of the maximum number of possible connections: 0.66 for the macaque (Markov et al., 2011) and 0.97 for the mouse (Gămănuţ et al., 2018). At such high values, and especially for the mouse, network specificity is achieved by the profiles of connection weights (Gămănuţ et al., 2018). The connectivity profile of a cortical area is the set of connections and their weights, which has been hypothesized to constrain its functional properties thereby reflecting its specialization (Bressler, 2004; Bressler and Menon, 2010; Markov et al., 2011).

Studies of existing, self-consistent tract-tracing datasets (Kennedy et al., 2013) reveal the action of a simple rule in both mouse and monkey, the so-called Exponential Distance Rule (EDR), which significantly constrains the structure of the interareal networks (Ercsey-Ravasz et al., 2013; Theodoni et al., 2020; Horvát et al., 2016; Markov et al., 2013a). The EDR expresses the empirical observation that axonal connection probability decays exponentially with projection length, *p(l) ~ e^−λl^*, where ⟨*l*⟩ = 1/*λ* is the average axonal projection length 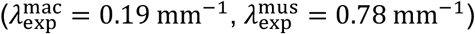. With the EDR, a one-parameter (*λ*), Maximum Entropy Principle based, generative model for interareal networks captures many binary, and some weighted features of the cortical network, including the frequency distribution of 3-motifs, global and local efficiencies, core-periphery structures, eigenvalue distributions and connection similarity profiles (Ercsey-Ravasz et al., 2013; Horvát et al., 2016; Theodoni et al., 2020; Song et al., 2014).

Interareal connections and the network structure are the evolutionary consequences of genetic pre-specification and interactions with the environment (Buckner and Krienen, 2013). Although there is network variability between individuals (Gămănuţ et al., 2018; Markov et al., 2014), one can speculate that there are universal features common to all individuals within species and across species (Goulas et al., 2019; Margulies et al., 2016; Mota et al., 2019). This is supported, for example, by the cross-species consistency of the aforementioned EDR (Horvát et al., 2016; Theodoni et al., 2020) and the similarity of topographical ordering of the functional areas on the cortical mantle (Krubitzer, 2009).

Here we refer to the above-mentioned universal features, as *architectural network invariants,* which we argue, imply predictability of networks. To study this issue in a more general and systematic fashion, we turn to data prediction and machine learning methods. We show that these techniques can be used to assess the degree of predictability of brain networks and are therefore also usable for network imputation, i.e., to predict missing network data. Naturally, the accuracy of imputation is determined by the degree of predictability inherent in the data. Moreover, we argue that predictability methods can also be *used* as tools to study structure-function relationships in these networks. Overall, these methods address the following questions: “Are certain parts of the network more predictable than others?”, “How much information do individual parts of the network carry about the network as a whole?”, “How well can missing connections be predicted?”, “How does heterogeneity in predictability relate to cortical function and behavioral features of the species?”, “How does predictability in an order (e.g., primate) compare to predictability in another (e.g., rodent)?” and “Can we use predictability as a guide for further experiments?”

Two key methodological aspects of our approach are to be emphasized. First, predictability is primarily an inherent property of the data itself and not of the algorithm used. Although the quality of prediction algorithms varies wildly, even the best algorithm cannot and should not “predict” information that is not there (for example, in the case of two pieces of mutually independent data A and B). Secondly, great care has to be taken in order to avoid overfitting, that is, fitting to noise in the data, as this leads to loss of generalization power and erroneous conclusions.

## Results

First, we introduce the datasets in our study then give a brief summary of the prediction methods used and how they are adapted to work with retrograde tract-tracing datasets. We then present our main results on network predictability using cross-validation both at the binary and weighted levels, along with a comparison of predictability between rodent and primate cortical networks.

### Data description

We rely on two retrograde tract-tracing datasets, one for the macaque (mac) and the other for the mouse (mus). Both are cortico-cortical connectivity databases created with consistent methodology, having most of the data published in (Gămănuţ et al., 2018) in the mouse and a more limited dataset based on 29 injection areas in a 91- area atlas in (Markov et al., 2014) on the macaque. The mouse dataset 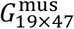 is a matrix of FLN values *FLNjj* for 19 injected target areas (*j* is a source, projecting into target *i*), in a 47-area parcellation. The current macaque dataset 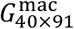 contains the original data for the 29 areas published in (Markov et al., 2014) and the weighted connections for an additional 11 areas, bringing the total of injected areas to 40 in a 91-area parcellation. Both datasets are provided in the Supplementary Information (SI). The full interareal networks (FIN), which are not available for either species, would be the matrices 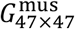 and 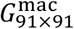, respectively. Additionally, our datasets contain all pairwise distances along estimated shortest paths avoiding anatomical obstacles, between the area barycenters, recorded in the matrices 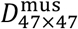 and 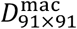, respectively, also provided in the SI.

Repeat injections across individuals allow an assessment of the consistency of the set of areas and their FLN values (Gămănuţ et al., 2018; Markov et al., 2011). Due to the high sensitivity of the tracers, every injection reveals all the areas that project into the injected target area and thus, the FLN matrix *G_T×N_* is a row submatrix of the FIN *G_N×N_*. That is, we either know the full row (corresponding to a target area) or not at all. This is illustrated in Figure 1A where, for simplicity, we order the rows such that the first *T* rows represent the targets, in the full *G_N×N_* matrix.

**Figure 1.**
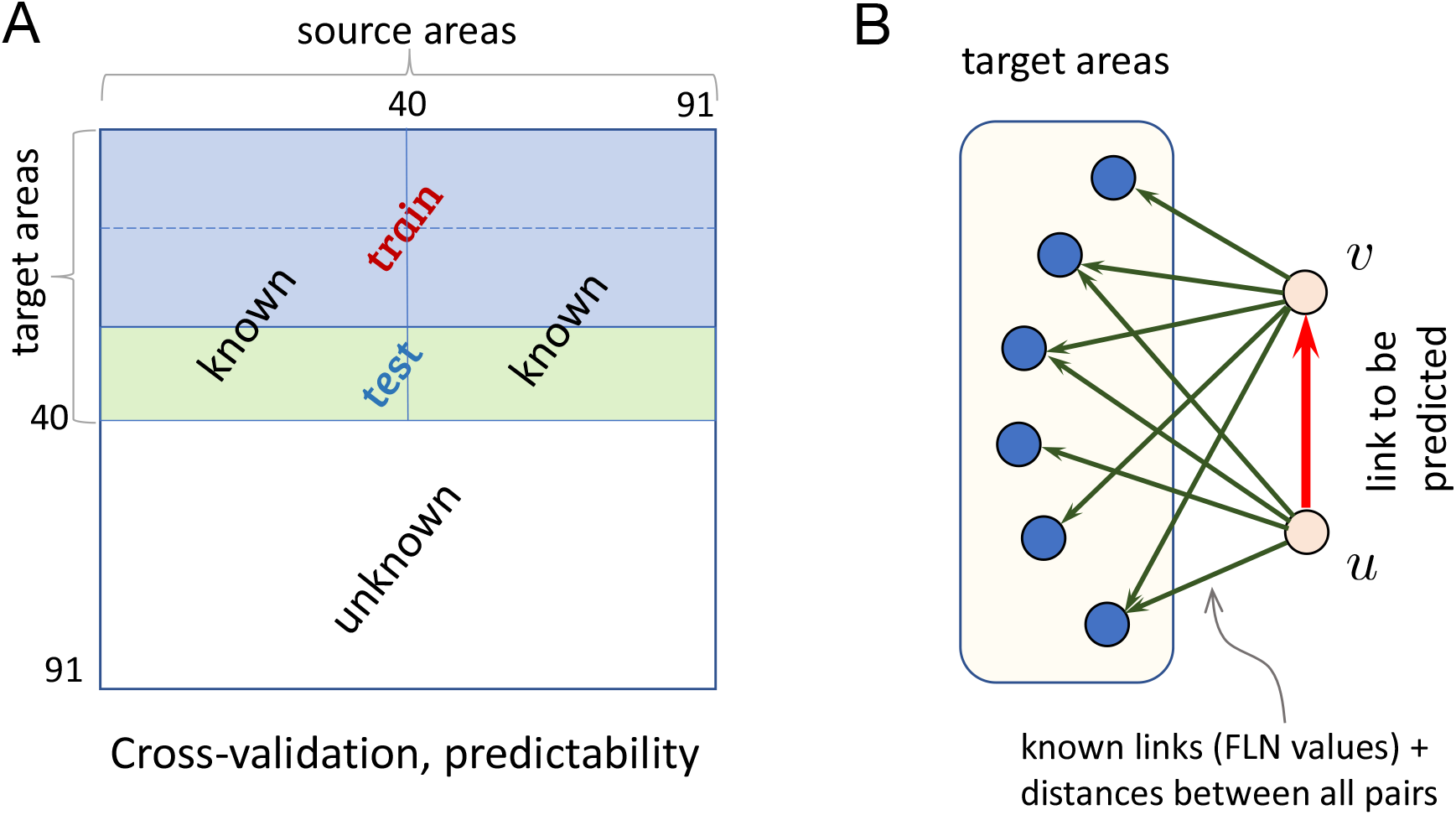
Schematics for link prediction with retrograde tract-tracing data. (A) *k*-fold crossvalidation setup for predictability (*k* = 3). (B) Links are predicted based on information (weights, distances) from the out-neighborhoods of its incident vertices.

#### Data preprocessing

In order to use the available input data, it needs to be organized in a format appropriate for the prediction algorithms (described below). We preprocess the FLN matrix by computing the base-10 logarithm of all its non-zero entries (Markov et al., 2013a) (values range in order of magnitude from 10^−7^ to 1) then shifting the values by adding 7 to them: *w_ij_* = 7 + log_10_(*FLN_ij_*). The zero entries are left as zeroes. The resulting matrix has values between 0 and 7 (in both species). The 0 entries correspond to non-links (i.e., non-connected node pairs) and elements on the main diagonal, non-zero entries to actual links. The macaque distance matrix range (0,58.2 mm), and the mouse (0,12 mm). For both species the distance feature matrix *D_f_* = 31 (*D*/max *D*) with values ranging from 0 to 31^1^.

### The link prediction framework

Link prediction refers to inferring links from observed network data (Liben-Nowell and Kleinberg, 2007; Clauset et al., 2008; Lü and Zhou, 2011). This can be done at the binary (predicting only if a link exists/1 or not/0) or weighted levels (predicting the associated weight). Binary level predictors also are known as classifiers, whereas weighted link predictors are essentially regressors. There are two main families of prediction methods for static networks, Classical Link (CL) predictors and Machine Learning (ML) predictors. CL predictors, used extensively in social networks, are classifiers that forecast links at the binary level based on either node neighborhood information (local) or path information (global). This information is formulated into a *predefined model* that generates a score *score(u,v)* for every ordered node pair *(u,v)*, which is then used to make the prediction. ML predictors can be used as classifiers (for binary prediction) or as regressors (for weighted prediction). They predict based on *learning* from samples with a given set of features. A feature is a vector of values (feature vector) quantifying what we know about the relationship of a node pair. We train an ML predictor in a supervised fashion, by providing the feature vectors computed for the node pairs in the training set and using the “ground truth” data about the pairs’ connectivity. The classifier then creates a model *autonomously* that best fits the given training set with the given feature vectors, which is then tested against the ground truth in the test data and the classifier’s performance is evaluated. Thus, the main difference between CL and ML is that we impose the model in CL, whereas it is learnt in ML. However, for both CL and ML, the information on which the prediction is based (scores and feature vectors) has to be computable *for all pairs in an identical fashion,* which limits the types of predictors that can be used for retrograde tracing datasets. In particular, for CL, path-based predictor models such as PageRank, Katz score, and Shortest Path score are effective when random links exist adjacent to the link (or non-link) to be predicted. However, in retrograde tracing datasets we are forced to select injected areas as the basis for predictions, but there are no paths into some of the vertices of the links to be predicted (i.e., the remaining areas that were not injected), thus excluding path-based predictors. For both CL and ML, we can only use information on out-going links, being the only type of information commonly available to all node pairs, see Figure 1B.

The performance of both classifiers (CL, ML) and regressors (ML) is evaluated using cross-validation techniques. This separates the available data with ground truth value into two sets: a training set and a test set. The former is used as input information for the predictor, which based on that, makes predictions for links in the test set, which is then compared to the ground truth. One of the two main approaches is the *k*-fold crossvalidation, used here, which splits the data into *k* equal parts, using in one iteration one of the parts for the test set and the other *k — 1* parts for training, then this is repeated for every combination of test/training split. Performance metrics are then averaged. To avoid correlations with any predetermined ordering of the input data we randomize the ordering of the target areas in the FLN matrices^2^ before splitting it into *k* parts. We then compute the corresponding averages over all these randomized realizations and all folds within. An alternative approach is Monte Carlo cross-validation, which we found gave very similar results to *k*-fold cross-validation.

For classifiers we use the standard receiver operating characteristic (ROC) curve and the corresponding single scalar metric, the area under the ROC curve (AUC) as performance metrics. The ROC shows the true positive rate (TPR) plotted against the false positive rate (FPR), obtained by moving the threshold value that distinguishes positive and negative predictions. A perfect classifier has 100% TPR and 0% FPR and the ROC curve fills the top left corner of the unit square; a random predictor has 50% TPR and 50% FPR with the ROC following the main diagonal of the unit square, whereas anything below the main diagonal implies an invalid predictor. The ROC curve also has a specific point that corresponds to the maximum prediction accuracy. Accuracy is defined as the number of correctly predicted links and non-links divided by the number of all predictions (ACC = (TP + TN) / (TP + TN + FP + FN)), where TP, TN, FP, and FN are the number of true positive, true negative, false positive, and false negative predictions, respectively. This point is determined numerically for each ROC curve, and this threshold is used to make the binary predictions during cross-validation. For weighted predictors there are no ROC curves. Instead, we use the mean absolute error (MAE) or the relative MAE (RMAE) between predicted and actual links weights (using RMSE, i.e., root-mean-square error gives very similar results).

Cross-validation helps to quantify not only how well a particular algorithm predicts the presence or absence of links but also to quantify the degree of predictability in the data, especially when comparing across ML algorithms, for both binary and weighted predictions. Note, predicting the connectivity status of node pairs for which there is no ground truth (imputation task), is only meaningful if the cross-validation results indicate significant predictability in the data. Here we present predictability results (crossvalidation) in both species using both CL and ML algorithms at binary and weighted levels. Link imputation will be presented in a subsequent publication.

## Network predictability in the macaque and mouse

### Binary link prediction

CL algorithms generate a score score (*u, v*) for every node pair (*u, v*) based on link predictor formulas that express various types of network information. These formulas, used typically in social networks, provide summations over nodes with incoming links from both *u* and *v*. Since retrograde tracing data such as the ones used here only reveal the incoming links to the target areas, the predictor formulas must be modified accordingly (shown in Materials and Methods).

In the case of ML classifiers, we need to specify the feature vectors. Figure 2 shows the macaque ROC curves for ML classifiers (solid lines) based on full information feature vectors, namely, feature vectors composed of both FLN values and distances. We have tested several other combinations of data for feature vectors and found the results to be invariably inferior to that based on full information (SI Figures S1-S4). ML classifiers other than those shown in Figure 2 have also been tested, such as DecisionTree, AdaBoost and NaïveBayes but overall had a performance inferior to those shown here. It is clear that with the exception of JA (modified Jaccard), the CL predictors do not perform as well as the four ML classifiers. The ML classifiers were tested against overfitting (SI Figures S6, S7 show the case of the MLP). They were also tested using different *k* values for the number of folds (SI Figure S8).

**Figure 2.**
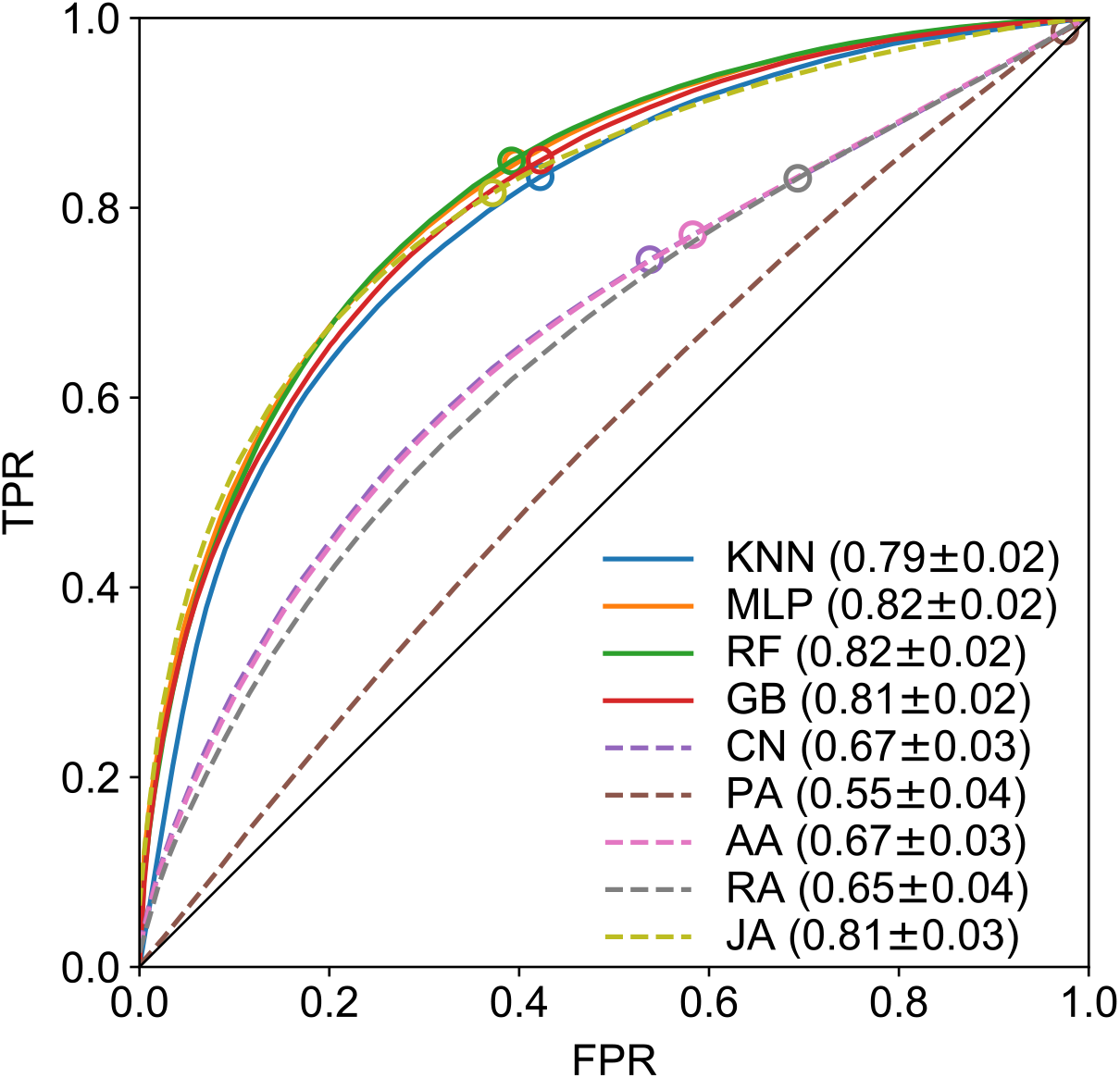
ROC curves for binary link prediction in the macaque. Dashed lines are from CL predictors: CN-common neighbors, PA-preferential attachment, AA-Adamic-Adar, RA-resource allocation, JA-Jaccard index. The continuous lines are from the four best ML classifiers, based on the full FLN-plus-distance feature vectors: KNN - K-nearest neighbors, MLP - multilayer perceptron and RF - random forest, GB - gradient boosting, using *k*-fold cross-validation, with *k* = 3. The markers indicate the location of the maximum accuracy thresholds.

The approximately 80% AUC obtained consistently by the top performing classifiers, indicates high predictability embedded in the macaque interareal network, suggesting the existence of architectural invariants and corresponding mechanisms (Figure 2). This analysis cannot be applied to the mouse dataset, (see the ROC curves in the SI Figure S9), due to its ultra-high density of 97%. This density causes a strong bias and prevents the calculation of a meaningful ROC curve, because the classifiers have only 3% true negatives to learn from, meaning that only weighted predictions can be made in the mouse brain, presented in the next section.

Figure 3 shows individual link prediction errors in the macaque data for all the links with a corresponding ground truth value (lighter colors correspond to smaller errors). A prediction (link existing/1 or not/0) was obtained for every *k*-fold run in all area-pairs *i*, averaged over 100 randomized *k*-fold run predictions, generating a prediction ⟨*y*_pred_(*i*)⟩.

**Figure 3.**
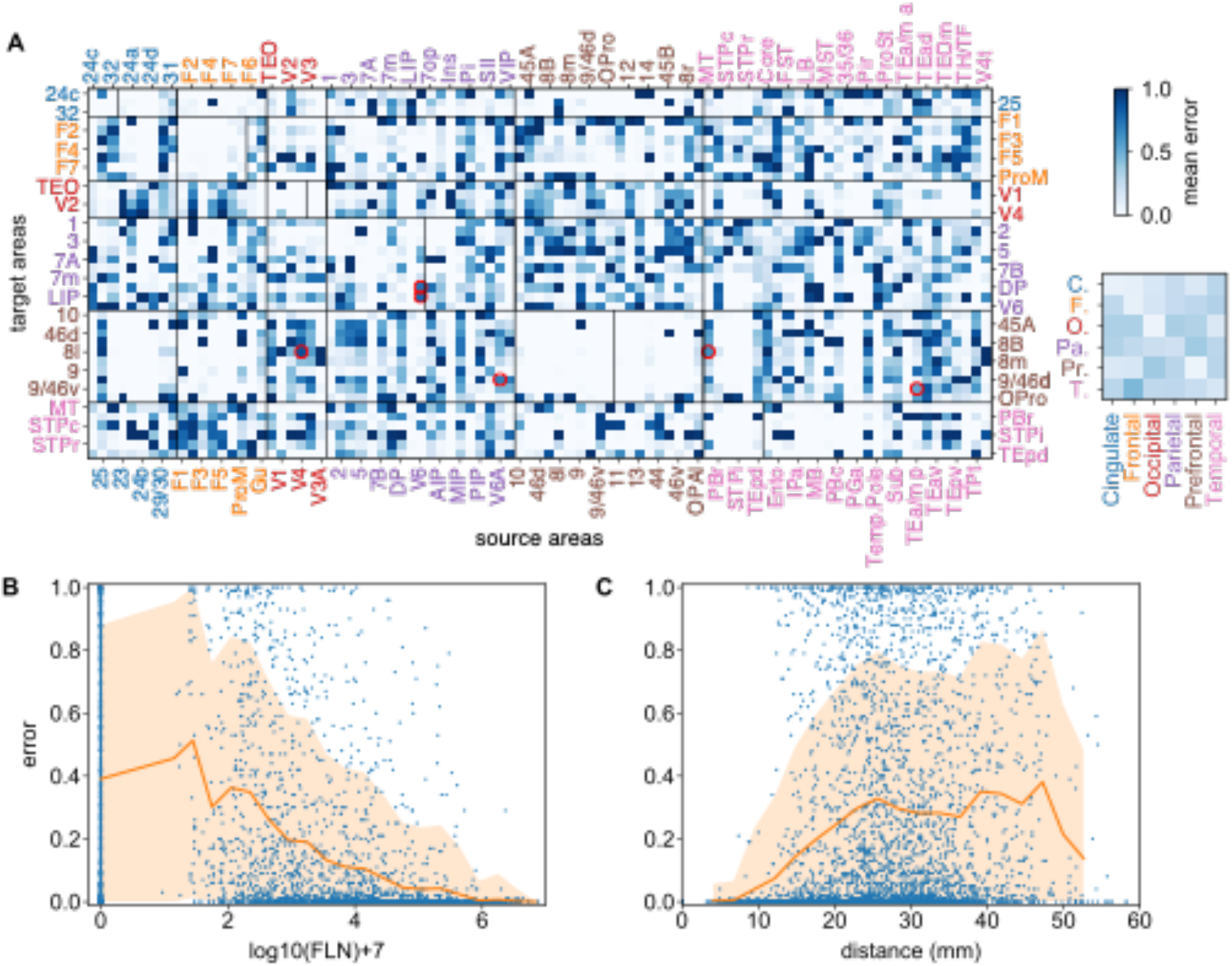
Binary prediction heterogeneity in the macaque brain. (A) Prediction error matrix for all known links (3-fold cross-validation) generated by GB. Vertical lines within the main diagonal boxes, separate targets (to the left of the line) from non-injected areas (to the right of the line). Red circles indicate strong links (with weights > 5) with high prediction errors. Along with their weights w and their errors ϵ, these are: V6 → DP (*w* = 5.3, ϵ =0.75), V6A→ 9/46d (*w* = 5.28, ϵ = 0.69), V6→LIP (*w* = 5.47, ϵ = 0.79), P4 → 8l (*w* = 5.11, ϵ = 0.87), MT → 8l (*w* = 5.33, ϵ = 0.62) and TEa/mp → 9/46v (*w* = 5.3, ϵ = 0.53)). Inset matrix shows inter-regional errors obtained by averaging errors within sub-matrices corresponding to cortical lobes. (B) Prediction errors as function of link weights and (C) as function of link projection distance. The vertical line in (B) at 0 are all the node pairs for which the prediction was *non-link,* while (C) contains all *links* and all *non-links.* The orange shaded areas in (B) and (C) represent one standard deviation from the average (orange line). The definition of error measure is given in the main text. Area abbreviations with corresponding area names and region assignments are provided in the SI Table S1.

The error is calculated via *error(i) = |y*_true_(*i*) — ⟨*y*_pred_(*i*)⟩|. where *y*_true_(*i*) ∈ {0,1} is the true binary link value. The inset in Figure 3A is a matrix of link prediction error heterogeneity by cortical brain regions. This shows that links from the frontal to temporal regions are less predictable (bottom row, second column), while links from frontal to cingulate (and prefrontal) are more predictable, etc. In addition, links within functional regions are more predictable than between regions (main diagonal of the small matrix), suggesting that predictability is possibly distance and thus weight dependent, since from EDR, we know that short/long connections are preponderantly strong/weak. Figure 3B,C show how prediction errors behave as a function of the link weights and distance demonstrating the action of a distance rule on predictability.

In order to disentangle the effect of distance/weight, we examined predictions based only on links of certain strengths: Strong only, *w_ij_* ≥ 5; Medium-&-Strong, *w_ij_* ≥ 3; Medium-&- Weak, *W_ij_* ≤ 5 and Weak only, *w_ij_* ≤ 3. The sizes of these weight groups are: 494 links for Strong, 1600 links for Strong-&-Medium, 3146 links for Medium-&-Weak and 2040 links for Weak. Figure 4 clearly shows that weak links are not predictable at the binary level (panel D) implying that the weak (thus long-range) links carry no information about each other. This is a significant observation that we revisit below, in our weighted prediction analysis as well. The maximum binary predictability is within the Strong-&-Medium group. The Strong group has a somewhat weaker predictability, possibly because that is the smallest set to learn from and the presence of some strong links with high unpredictability (red circles in Figure 3A). One of them, V4 → 8l is part of a strong loop, discussed in the literature (Markov et al., 2013b, 2013a; Vezoli et al., 2021). Note that these are the links with the highest prediction errors within the Strong group, *only.*

**Figure 4.**
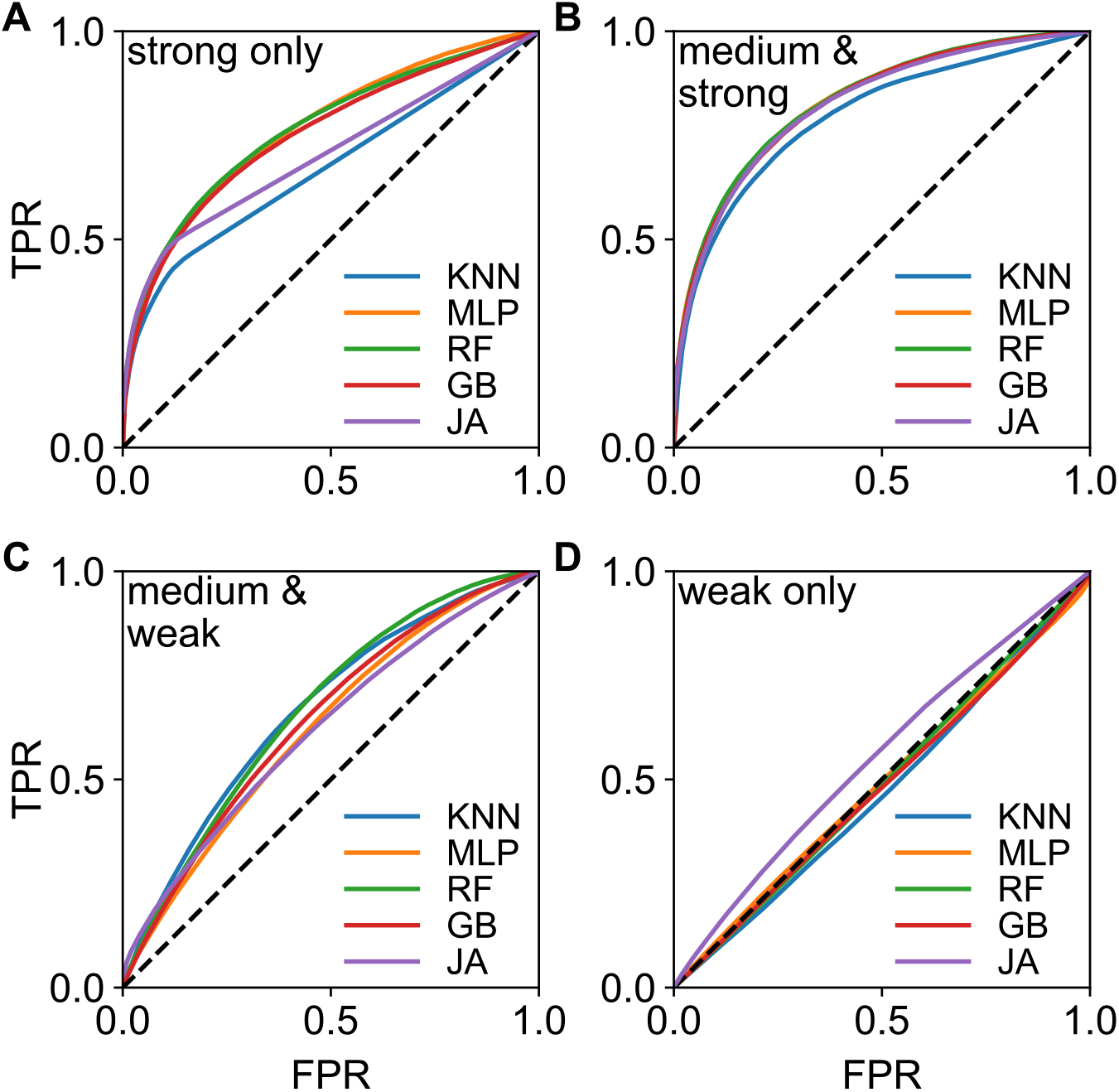
Binary predictability as function of link weights. Predictability from only (A) Strong links *w_ij_* ≥ 5 (494 links), (B) Strong-&-Medium *w_ij_* ≥ 3 (1600 links), (C) Medium-&-Weak *_ij_* ≤ 5 (3146 links) and (D) Weak links *w_ij_* ≤ 3 (2040 links). The AUC values and errors in (A) KNN (0.67 ±0.02), MLP (0.77 ± 0.03), RF (0.77 ± 0.02), GB (0.75 ± 0.02), JA (0.70 ± 0.02); in (B) KNN (0.79 ±0.02), MLP (0.83 ± 0.02), RF (0.83 ± 0.02), GB (0.83 ± 0.02), JA (0.82 ± 0.02); in (C) KNN (0.67 ±0.02), MLP (0.62 ± 0.04), RF (0.67 ± 0.03), GB (0.64 ± 0.03), JA (0.62 ± 0.04); in (D) KNN (0.47 ±0.03), MLP (0.49 ± 0.05), RF (0.49 ± 0.03), GB (0.48 ± 0.03), JA (0.55 ± 0.02).

### Weighted link prediction and comparisons between mouse and macaque

In order to predict the link weights, we need to turn to supervised regression methods. This excludes CL algorithms as they are designed uniquely for binary link predictions. Since all our ML classifiers are available as regression algorithms as well, they can be readily used for weighted link prediction. The same feature vectors as for binary classifiers are used but the ground truth now is the actual link weight, *w*_true_. In terms of evaluating the performance and the amount of predictability inherent in the network we employ the *k*-fold cross-validation scheme as previously, but the performance metric has to be modified (there are no ROC curves in weighted link prediction). Here we could use the mean absolute error (MAE) obtained as the absolute value of the difference between the predicted and the actual weight |Δ*w*| = |*w*_pred_ − *w*_true_|, averaged over the 100 *k*-fold predictions. Since FLN values vary over orders of magnitude, the MAE of a weak link is not easily comparable to that of a strong link. In order to take this into account, we employ the relative MAE (RMAE), which is the MAE divided by the ground truth strength of the predicted link, |Δ*w*|/*w*_true_. Thus, the RMAE value is the fraction of the link weight that is not predicted. For example, an RMAE of 0.2 means that 80%of the link weight *w* was predicted and 20%was not. An RMAE of 2 reflects an error of 200%compared to the true link strength. As for the binary prediction, comparing the performance of several classifiers, GB, KNN, MLP, RF come out as the four top predictors.

These regressors work by minimizing a cost function (such as the root-mean-square error RMSE) over the training set, when finding the best-fitting model, which in turn, is used to predict the test set. Analysis of prediction residuals provides both an efficient test of the capacity of the predictor to capture the signal part of the data as well as a means of ranking performance. This analysis shows that GB performs somewhat better compared to RF, MLP or KNN. SI Figure S10 shows the results from the analysis of the prediction residuals for the GB algorithm. A featureless scatter plot of the residuals vs. predicted values as shown in SI Figure S10C indicates that the signal portion of the data has been well learned by the predictor.

For simplicity, in the following we show predictions based only on GB. Figure 5A,B shows the prediction error (RMAE) matrices for both the macaque and mouse. Note the strong similarity of the patterns between Figure 5A and Figure 4A for the macaque. At the weighted level as well, some of the links are more predictable than others. The matrices at the regional level, presented in Figure 5C,D also show heterogeneity: for example, across species, temporal to occipital are highly predictable, whereas occipital to frontal are less so. Globally, the mouse network appears more predictable than the macaque (overall lighter matrices for the mouse). This is further demonstrated in Figure 6 where we plot RMAE values as function of link weight as well as a function of link projection lengths (distance). While in both species, weaker links are harder to predict, comparing (A) to (C) we see that the medium-to-strong links are much more predictable in the mouse than in the macaque, but the situation is reversed for the weakest links. Similarly, long-range links are harder to predict in both species than shorter ones. Overall, weighted links are more predictable in the mouse than in macaque.

**Figure 5.**
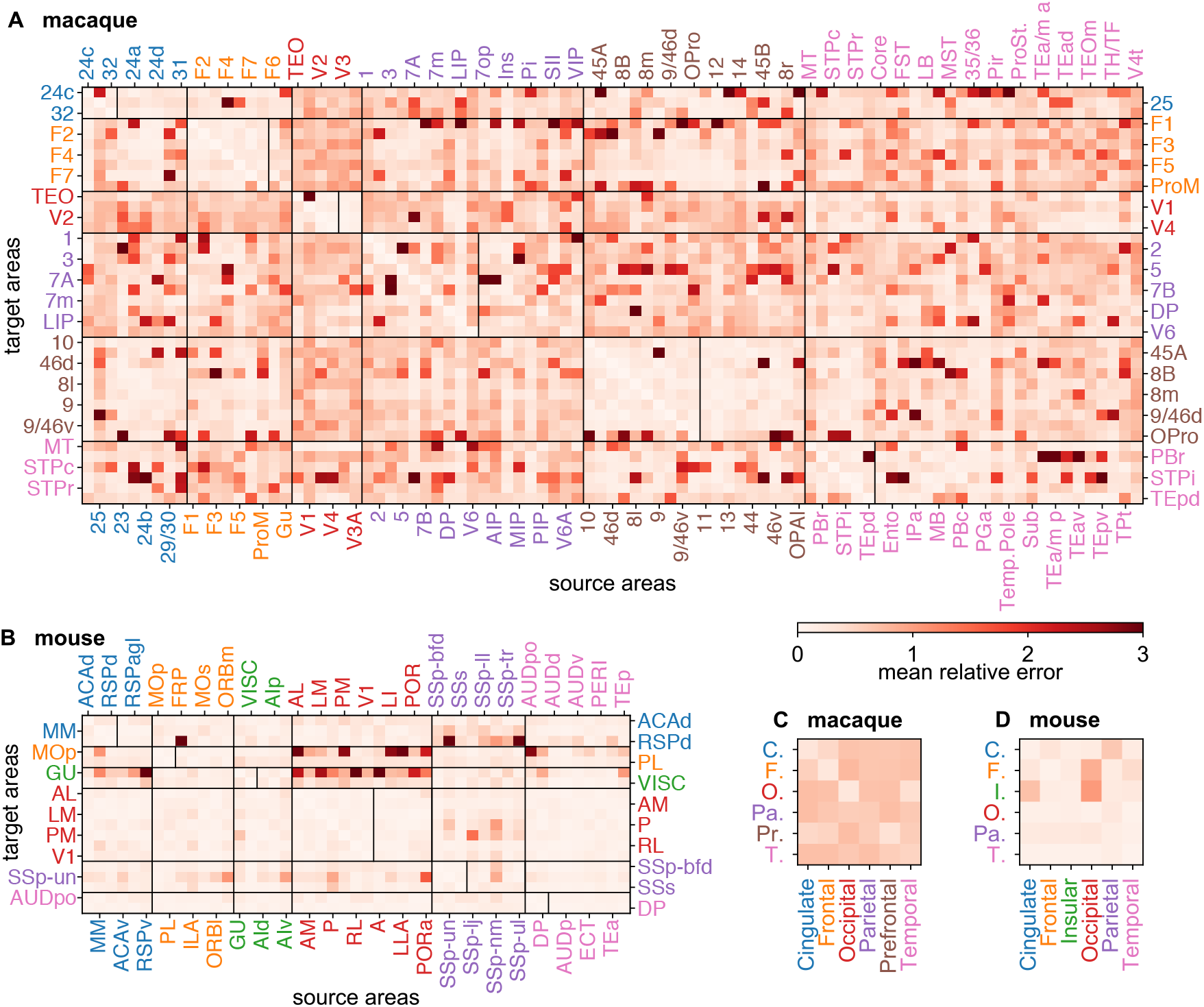
Prediction error heterogeneity for link weights. (A) Weight prediction error (defined as relative mean absolute error, RMAE) matrix for all known links with 3-fold cross-validation, in the macaque, generated by GB and (B) in the mouse. The vertical lines within the main diagonal boxes, separate targets (to the left of the line) from non-injected areas (to the right of the line). (C) inter-regional error matrix for the macaque (averaged from the matrix in (A)) and (D) for the mouse (averaged from the matrix in (B)). For non-links, the RMAE was calculated using the lowest statistically acceptable FLN value of 8 × 10^−7^ for the ground truth value (corresponding to a weight of *w* = 0.9). Area abbreviations with corresponding area names and region assignments are provided in the SI Table S2.

**Figure 6.**
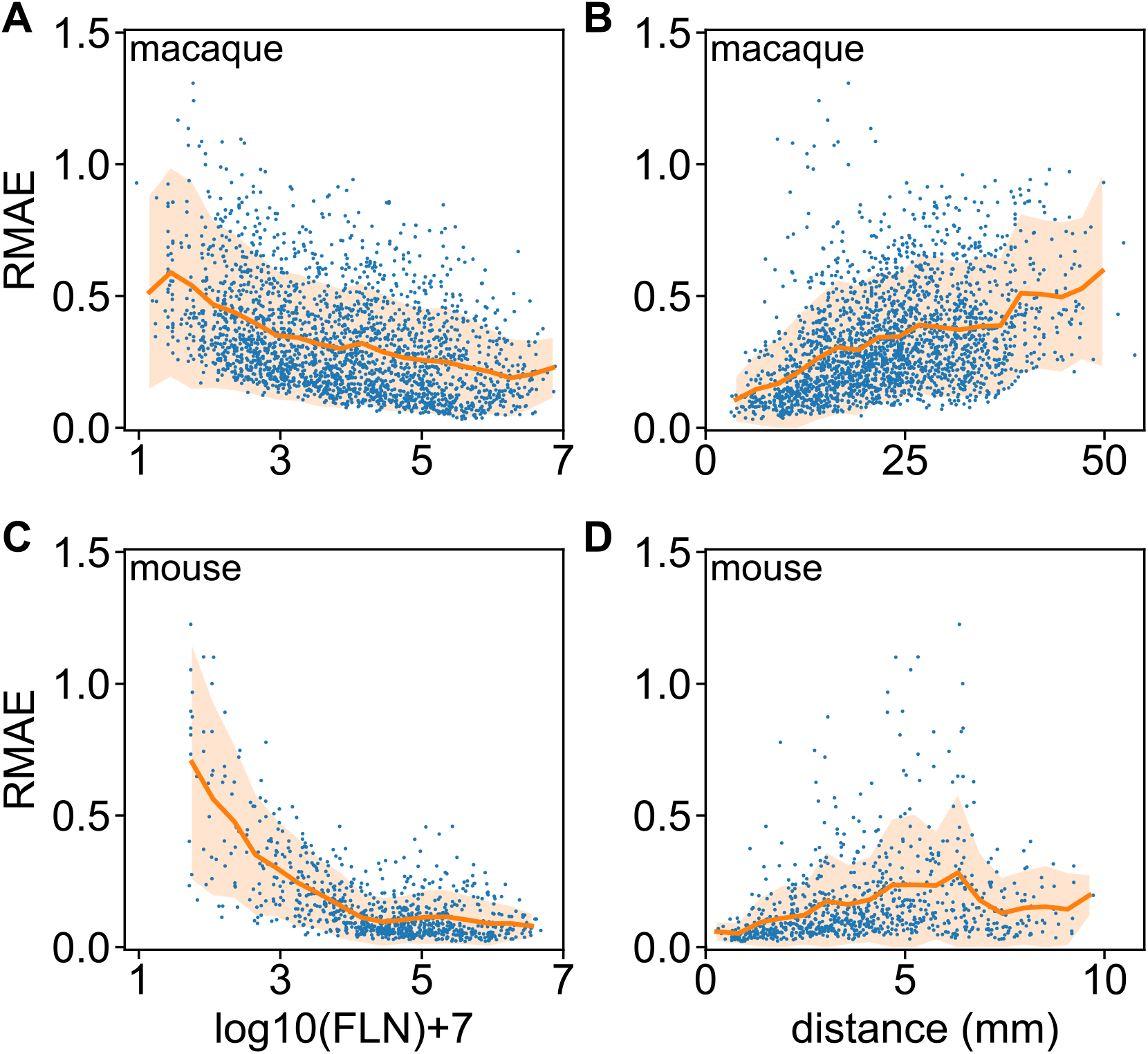
Weighted prediction errors as function of link strength and distance, using the prediction data from. Figure 5. (A) Relative mean absolute error RMAE vs link weight and (B) vs projection distance, in the macaque for every predicted link. (C) same as (A) and (D) same as (B), for the mouse. The continuous line is the mean value, the orange shaded area corresponds to one standard deviation. Panels do not contain data for no connections.

We quantify global link predictability, and by link weight classes in Table 1, for both species. Predictions (3-fold cross-validation) were made on the full dataset (including links with non-zero weight and also non-links) using the GB algorithm and errors computed and averaged within the corresponding groups. The RMAE values in Table 1 show that weak links are not predicted in either species, whereas the stronger links are better predicted in both species. The stronger links are in general two-fold more predictable in the mouse than in the macaque. The non-links, however, are better predicted in the macaque, likely due to the fact that there are only 3%non-links in the mouse dataset. Since the larger errors are associated with the non-links, we performed the predictability analysis also on a reduced dataset, with only actual links included (non-links excluded). That is, we trained the ML algorithms only on the portion of the matrix with non-zero link-weights. The predictability results are shown in SI Table S3. Except for the weak links, predictability improved in general, with mouse links being overall 1.5 times more predictable than the macaque ones.

**Table 1.** Prediction errors by link weight. MAE: Mean Absolute Error |Δ*w*|= |*w*_pred_ − w_true_|, RMAE: Relative Mean Absolute Error |Δ*w*|/*w*_true_. For “non-links” only, for the relative error, we used the estimated experimental lower cutoff value of *w*_true_= *w*_cut_= 0.9, corresponding to an *FLN* = 8 × 10^7^.

| Non-links included | Macaque |  | Mouse |  | Mac/Mus |
| --- | --- | --- | --- | --- | --- |
|  | MAE | RMAE | MAE | RMAE | RMAE ratio |
| Weak ($w_{\text{cut}} < w < 3$) | 1.016 | 0.441 | 1.032 | 0.446 | 0.989 |
| Weak-&-Medium ($w_{\text{cut}} < w < 5$) | 1.134 | 0.358 | 0.647 | 0.196 | 1.827 |
| Medium-&-Strong ($w > 3$) | 1.227 | 0.282 | 0.565 | 0.127 | 2.220 |
| Strong ($w > 5$) | 1.272 | 0.228 | 0.569 | 0.102 | 2.235 |
| All links ($w > w_{\text{cut}}$) | 1.164 | 0.330 | 0.622 | 0.166 | 1.988 |
| Non-links ($w \leq w_{\text{cut}}$) | 1.382 | 1.039 | 2.911 | 2.288 | 0.454 |
| Both links and non-links | 1.246 | 0.591 | 0.683 | 0.222 | 2.662 |

### Scaling of predictability with input data, leave-one-out analysis

Another important issue is the scaling of predictability with the amount of input, i.e., the amount of data used for training. To investigate this question, we consider a random subset of *m* areas from all the targets, leave one *target area* out (of this set of *m*), then make a prediction based on the rest for all the out-links of this one area. We then repeat this with every member of this subset, obtaining predictions for all of them. These are then compared with the ground truth and the relative error is calculated. We call this the *internal relative error* (internal to the selected subset). We then repeat this random selection of *m* subsets 500 times and average the obtained relative errors for all the targets, shown in Figure 7. An interesting conclusion from Figure 7 is that the ML predictors are able to learn the structure in the data fairly quickly for the medium to strong links, and the improvement after that is relatively small, although more significant for the weak links (note the log-scale on the y-axis). Another way of studying the scaling of errors with input data size is described in the caption of SI Figure S12, which shows prediction errors for areas external to the *m* selected. The two plots are not significantly different, leading to the same conclusion.

**Figure 7.**
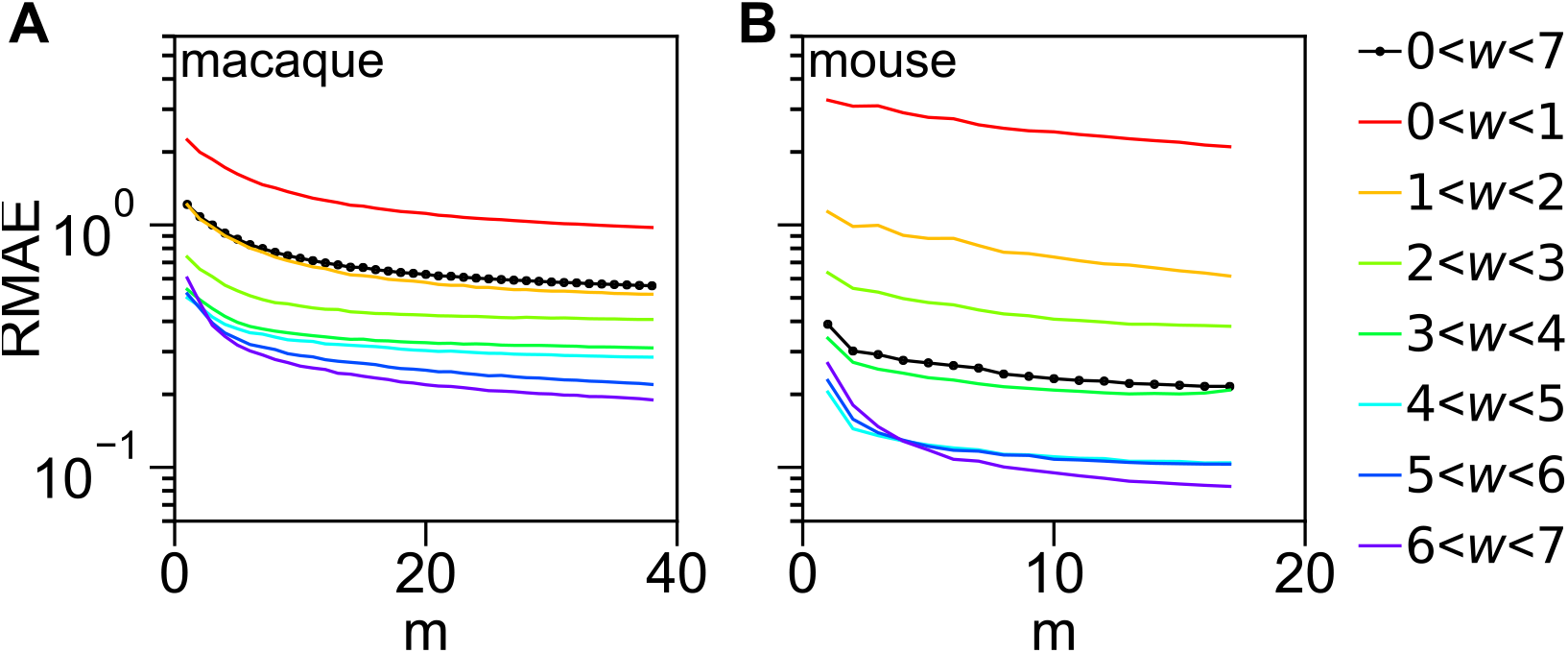
Scaling of prediction errors as function of input data size in a leave-one-out analysis. The relative mean prediction errors RMAE (of weights) are computed for areas internal to a set of *m* targets for both macaque (A) and mouse (B), then plotted as function of m, see main text for description. The errors are separated by link weight class. Note the logarithmic scale on the y-axis.

## Discussion

Using machine learning methods, we demonstrated that the network of the mammalian cortex contains significant structural predictability, suggesting that the formation and evolution of the cortex is to a good extent rule based, at the mesoscale level. This further motivates the search for universal mechanisms of brain network formation and evolution, within the mammalian order.

The literature on link prediction in the brain is fairly limited. To the best of our knowledge, the earliest link prediction effort in the context of brain neuronal networks goes back to a 1998 paper by Jouve et al. (Jouve et al., 1998), which uses the frequency of directed transitive triples to predict missing links at the binary level (existing or not), in an early dataset on the macaque visual system (Felleman and Van Essen, 1991). The next brain link prediction papers appear almost a decade later, which incorporate additional topological and spatial features (Costa et al., 2007; Nepusz et al., 2008), both based on the CoCoMac database (Kötter, 2004), with the latter using a stochastic graph fitting method to handle the uncertainties in the data. Several other publications followed these papers (Hoff, 2009; Cannistraci et al., 2013; Hinne et al., 2017; Røge et al., 2017; Chen et al., 2020; Shen et al., 2019), but all (including the earliest three) are based on preconceived network models whose parameters are fitted to the data, and then used to make predictions (usually at a binary level) on missing links. These network models quantify the belief that the existence or absence of a link is largely determined by some summary network statistics on the existing data. One problem with this approach is that it imposes specific relationships that the modeler believes to be relevant. Another is that the summary statistics are obtained on an incomplete dataset, which inherently biases these statistics, a bias which is then built into the prediction. A further bias present in almost all the previous link prediction papers is that they are based on network models from the field of social networks. However, brain neuronal networks are quite different from social networks in many aspects and thus social networks-based models would have limited practical applicability in the brain. Here we compared the performance of the social science inspired, model-imposed link predictors (CL) with machine learning based methods (ML) that learn the structure from the data, without imposing specific models or assumptions. Our results show that the latter approach achieves significantly better predictions than the model-based predictors. Another reason for the poor performance of most CL predictors is the fact that the CL formulas use only a single weight value and not multivariate information about a link (such as weights plus distance) efficiently, unlike ML algorithms (using only distances for CL, gives worse performance, see the SI Figure S5). The Jaccard coefficient is the only successful CL predictor because its formula happens to agree with a property of the link weight distributions in the brain. More precisely, it is due to the fact that the formula for the Jaccard index correlates with the triangle inequality, which holds for spatial networks and that also happens to be respected by the link weights of the brain, due to the action of the EDR: if areas A and B are close to each other (strong link) and area C is far from A (weak link), then C will also be far from B (weak link), mimicked by the Jaccard index as well.

Another significant issue affecting the reliability of predictions is the quality of the dataset. While the CoCoMac database is one of the largest connectomics databases, it is also a collation of results from independent studies using different approaches under different conditions, with significant inconsistencies (Bezgin et al., 2012). Moreover, the CoCoMac database does not distinguish absent links from links of unknown status, which is a major source of errors in network modeling (Kennedy et al., 2013). The results generated from the two retrograde tract-tracing datasets in non-human primates (macaque) and rodent (mouse), both obtained with consistent empirical methodology, allow for interspecies comparisons (Horvát et al., 2016) of the statistical network properties, using the edge-complete portions of the datasets. The ML predictions show that weak/long-range links in general are not predictable in either species, and that these links have little to no information about each other, at least in terms of link weights and projection distance. Moreover, they also show that overall, the mouse brain has a more predictable structure than the macaque (roughly by a factor of two in terms of errors). However, it is somewhat more difficult to predict the weakest connections in the mouse, than in the macaque (compare panels A and C in Figure 6). Accordingly, one could speculate that the long-range connections have less specificity in the mouse brain than in the macaque. It is, however, important to note that these predictability measures are all based on the features of link weights and projection distances. Including additional, biologically relevant features such as cell types could lead to a refinement of the predictability analysis presented here.

Finally, we recall that the EDR model (Ercsey-Ravasz et al., 2013; Markov et al., 2013a; Horvát et al., 2016), mentioned in the introduction, captures many features of the cortical networks in both species. One may ask, what is the amount of predictability in the EDR model networks themselves, using the same distance matrices as in the data, and the corresponding, empirically obtained *λ* decay rates? We find that the top predictors achieve a slightly better performance on the EDR model networks (an AUC of 0.86, see SI Figure S11) than on the experimental connectivity data (an AUC of 0.82, see Figure 2). The improved performance in the EDR network is expected, given that these networks are, by definition, rule-based, with some level of randomness included (Ercsey-Ravasz et al., 2013).

Machine learning methods may also be used as a guide to future neuroanatomical experiments. For example, if all predictors consistently suggest the existence or absence of a link where the data indicates the opposite, it may prompt the revision of the empirical data. Prediction results could also propose optimal injection sites based on the expected surprise that the data could reveal from such injections. Targets that generate in-links that are highly similar to the existing data do not add much novelty to the dataset, but areas with large deviations from the average link predictions may contain significant information about the specificity of the incoming links into that target. These could correspond, for example, to the appearance of a novel information processing modality in the brain, reflecting a significant evolutionary branching event in the history of the species.

## Materials and Methods

### Software packages

For this work we used Python 3.7 and SciKit-Learn version 0.20.2. The computation of the ML and CL predictors, cross-validation, and analysis of the results were implemented in a Python. General calculations and plotting functions are utilizing the standard packages of NumPy and Matplotlib.

### Classical Link predictor formulas

Since we do not have incoming links except for injected areas, we need to modify slightly the predictor formulas as shown in Table 2.

**Table 2.**
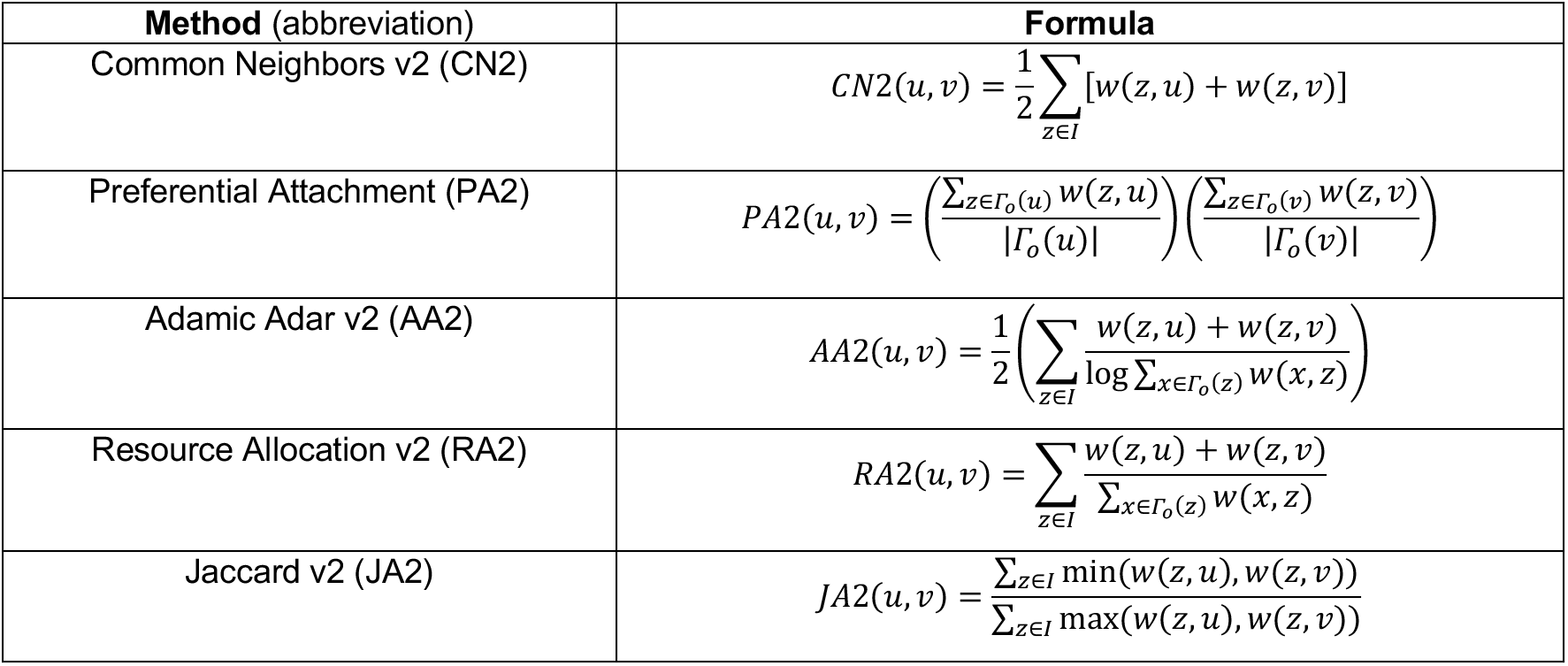
Classical, neighborhood-based link predictors for directed and weighted networks. The formulas have been adapted to be based on the out-link neighborhood information of the endpoints (*u, v*) of the directed link to be predicted. Each formula provides a prediction score *s(u, v)* for that directed link. Here *I* denotes the set of all target (injected) areas and Γ_0_(*u*) denotes the neighbors of *u*, including itself.

### Machine learning classifiers and predictors

All the classifiers used are implemented in the Python package scikit-learn; “defaults” refer to those parameters provided in version 0.20.2 of the library. We list the other parameters used for each classifier below.

- K-Nearest Neighbors (KNN): n_neighbors = 5, leaf = 30
- Decision Tree (DT): defaults
- Random Forest (RF): n_estimators = 200, criterion =‘gini’
- Multi-Layer Perceptron (MLP): hidden layer size: 100, convergence error tolerance: 10^-6^, max iterations: 20
- Gradient Boosting (GB): n_estimators = 100 (default), which is the number of boosting stages to perform. GB is robust to over-fitting and larger values typically yield better performance. max_depth = 7 (not default). This is the maximum depth of the individual regression estimators. It limits the number of vertices in the tree.
- AdaBoost (ADA): defaults
- Naïve Bayes (NBA): defaults

### Feature vectors

Here we summarize the feature vectors that we used to train and test the classifiers. In each feature function in Table 3, the link in question is *(u,v); A* denotes the weight matrix; *D* denotes the distance matrix; *d(x)* denotes the outdegree of node *x* in *I*; and *I* denotes the set of injected areas (nodes). Notice that the feature vectors have various lengths, as some provide more information than others.

**Table 3.** Machine learning feature functions used to train our classifiers.

| Feature | Formula |
| --- | --- |
| weighted_common_neighbors | $\sum_{i \in I} [A(i, u) + A(i, v)]$ |
| degree_plus_distance | $\{d(u), d(v), D(u, v)\}$ |
| adjacency | $\{A(i, u) > 0, A(i, v) > 0 \vert \forall i \in I\}$ |
| outdistance_source | $\{D(i, u) \vert \forall i \in I\}$ |
| outdistance_target | $\{D(i, v) \vert \forall i \in I\}$ |
| outdistance | $\{D(i, u), D(i, v) \vert \forall i \in I\}$ |
| fln | $\{A(i, u), A(i, v) \vert \forall i \in I\}$ |
| fln_plus_distance | $\{A(i, u), A(i, v) \vert \forall i \in I\} \cup \{D(u, v)\}$ |

## Acknowledgments

This work was supported by the National Science Foundation (NSF) grant IIS-1724297 (to Z.T. and H.K.), ANR-17-FLAG-ERA-HBP-CORTICITY (to H.K., M.E.-R., and Z.T.), ANR-19-CE37– 0025-DUAL_STREAMS (to K.K.) and by the CNCS-UEFISCDI grant COFUND-FLAGERA 2-CORTICTY (to M.E-R).

## Footnotes

1 This value gives a good resolution on the distance range, but other similar values can also be used.

2 The training of ML predictors can be sensitive to the order in which the training data is supplied.

