## Supplemental file for "Predictability of cortico-cortical connections in the mammalian brain"

**This PDF file includes:**

Figures S1 to S12
Tables S1 to S3
SI References

### Supplementary Figures

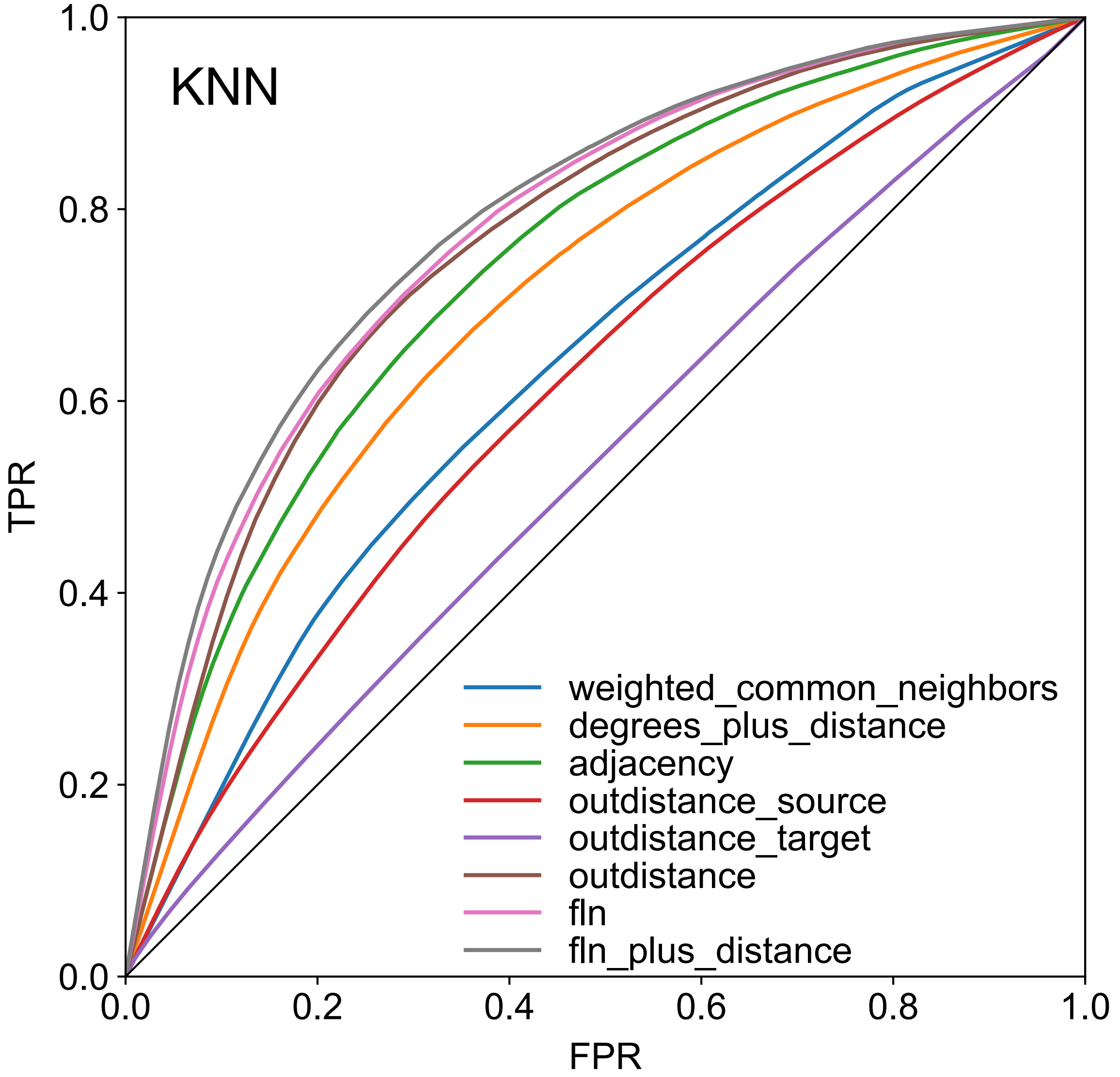


**Fig. S1 ROC curves for binary link prediction with various ML features in the macaque, using the KNN classifier.** Using 3-fold cross-validation, averaged over 100 samples.

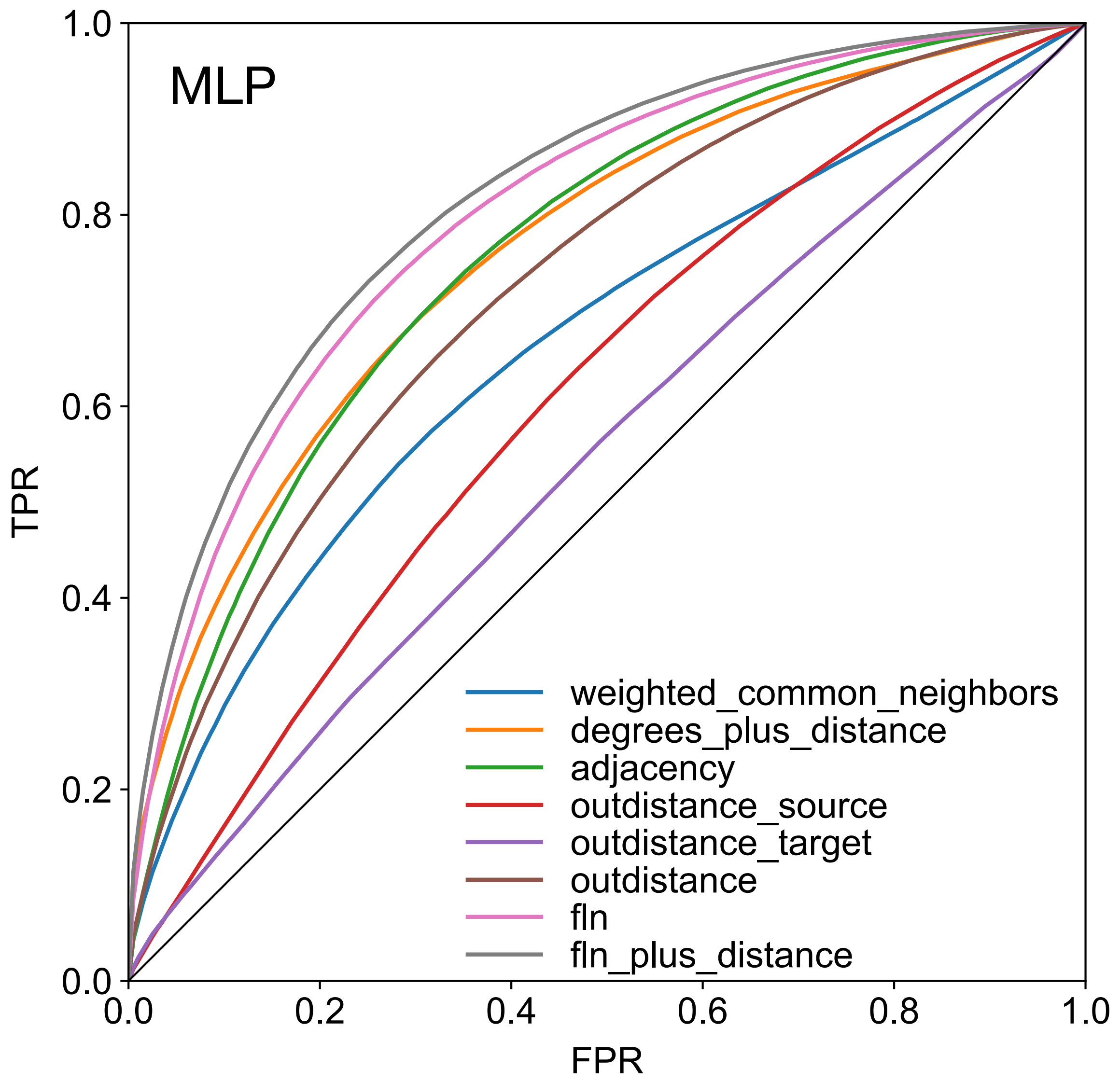


**Fig. S2 ROC curves for binary link prediction with various ML features in the macaque, using the MLP classifier.** Using 3-fold cross-validation, averaged over 100 samples.

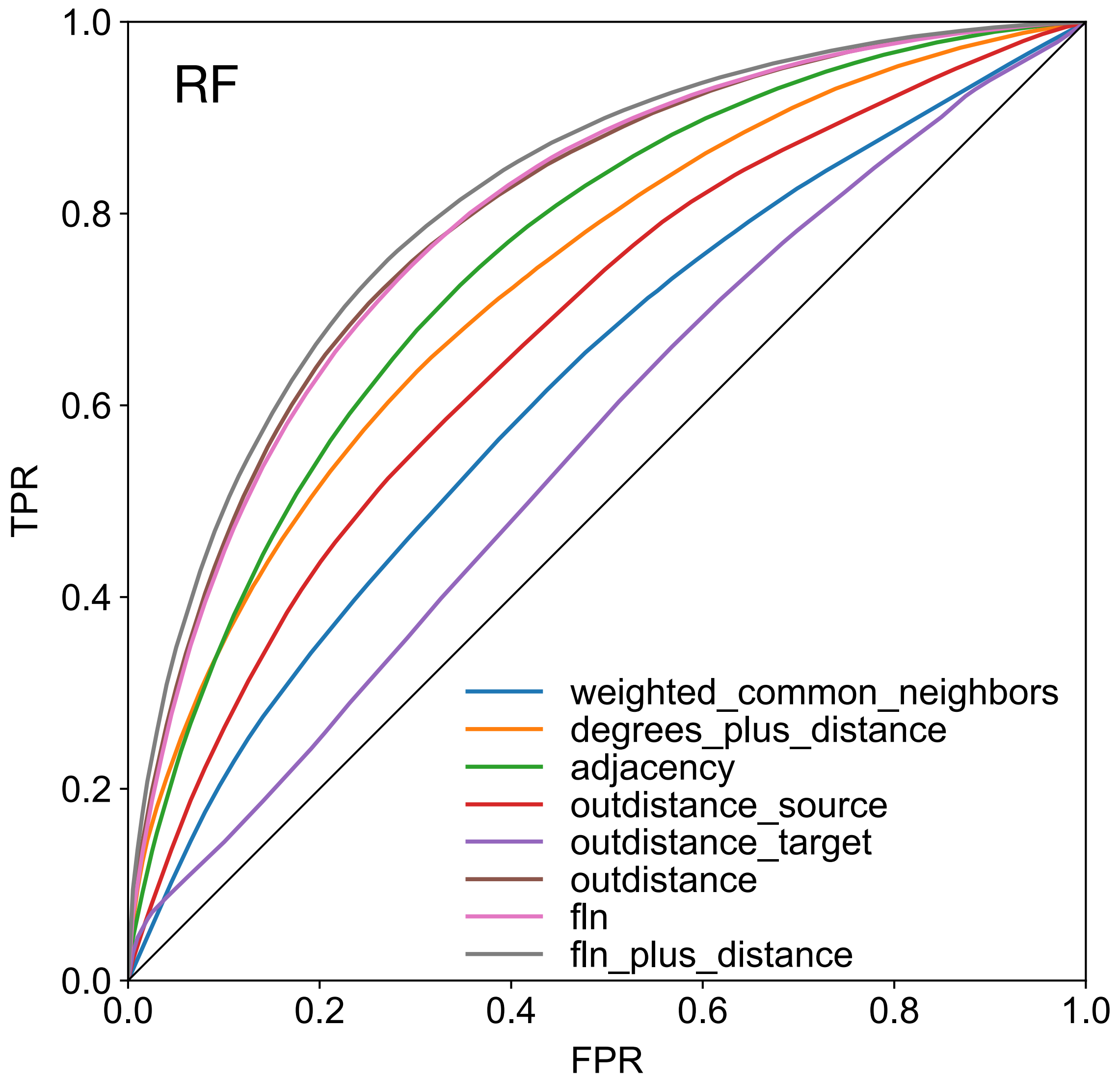


**Fig. S3 ROC curves for binary link prediction with various ML features in the macaque, using the RF classifier.** Using 3-fold cross-validation, averaged over 100 samples.

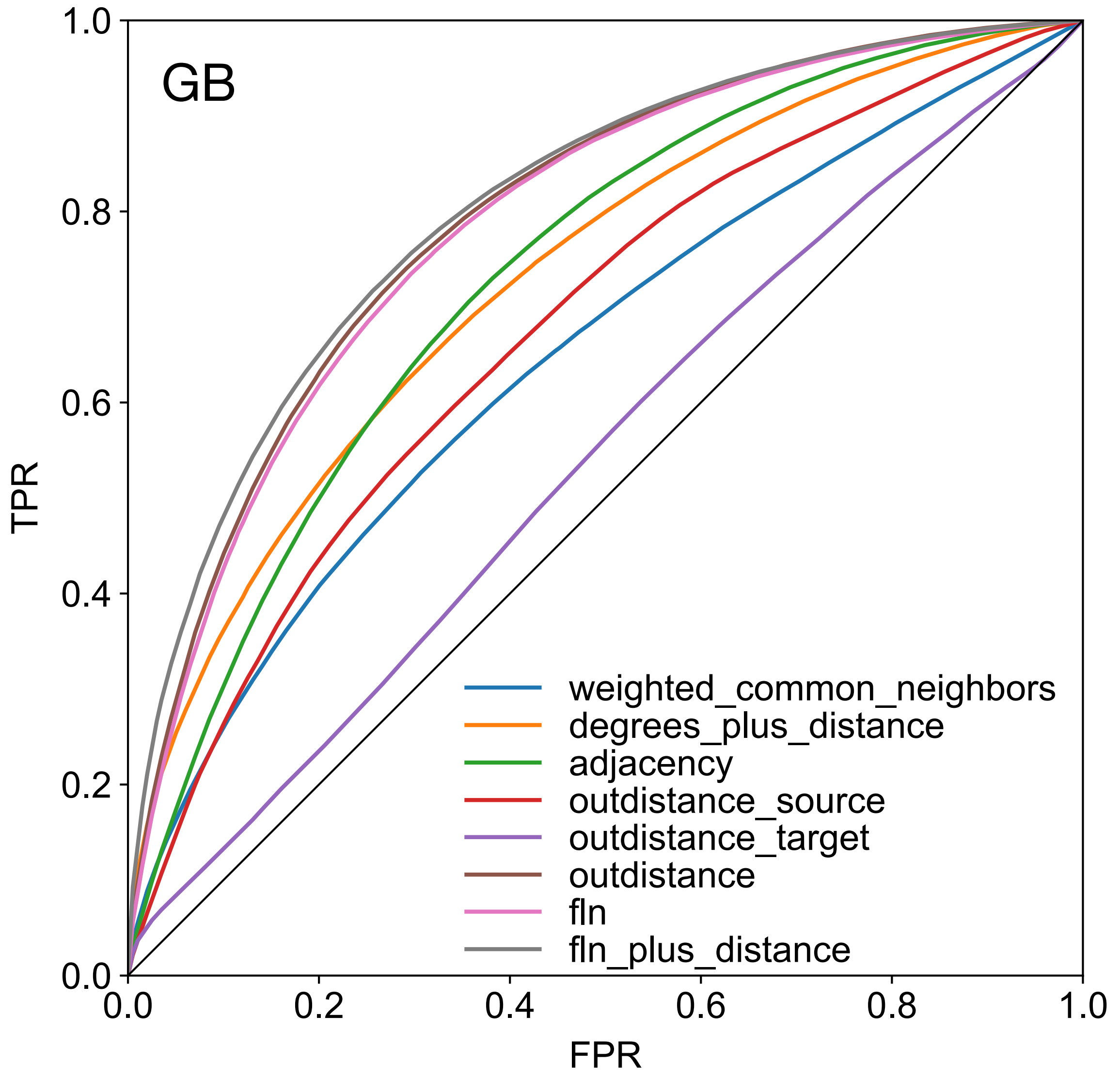


**Fig. S4 ROC curves for binary link prediction with various ML features in the macaque, using the GB classifier.** Using 3-fold cross-validation, averaged over 100 samples.

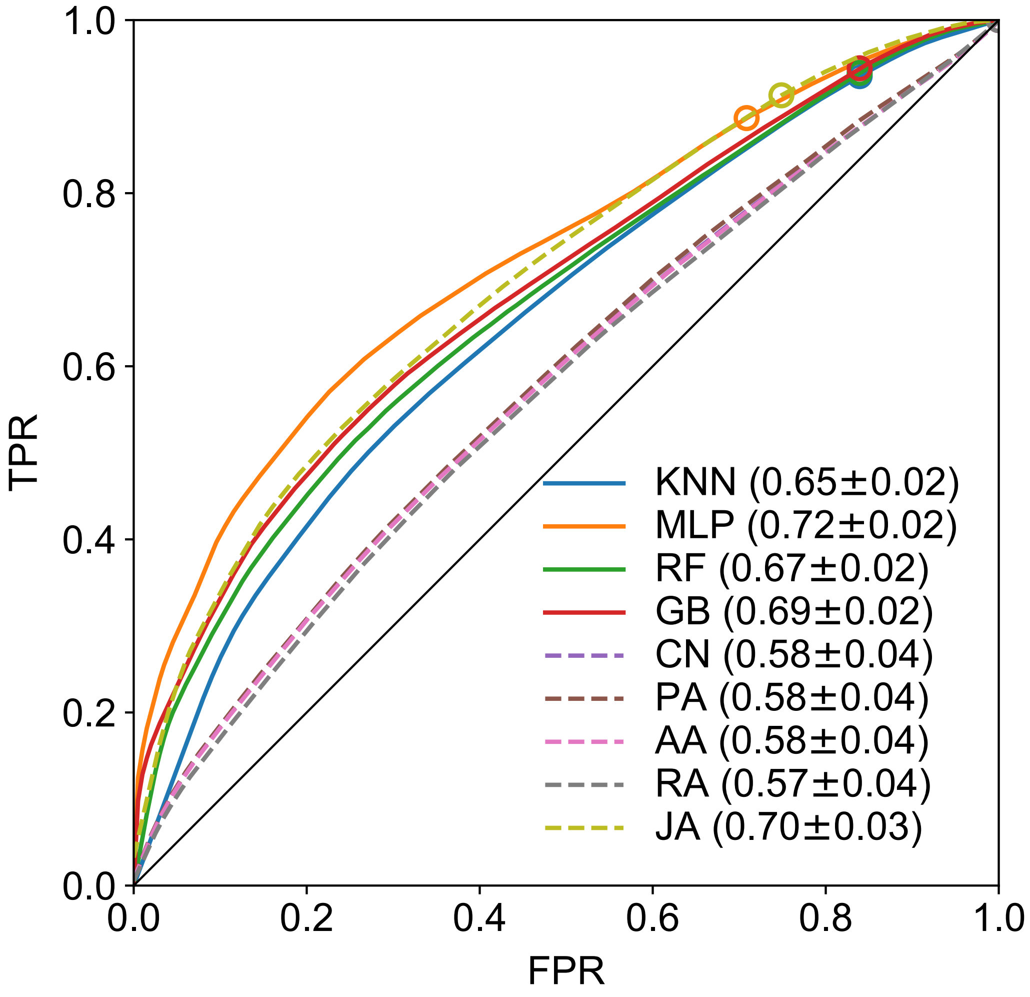


**Fig. S5 ROC curves for binary link prediction in the macaque, using only the distance feature and both ML and CL algorithms.** 3-fold cross-validation, averaged over 100 samples.

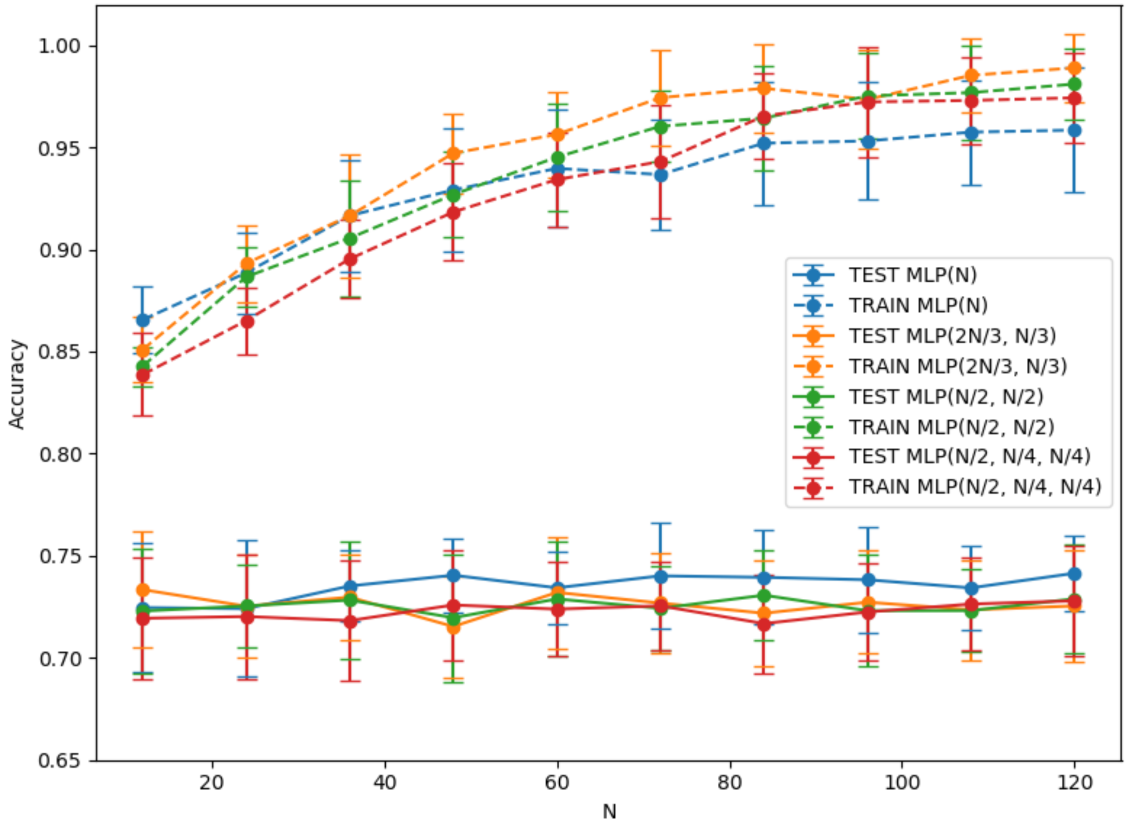


**Fig. S6 Testing overfitting of the MLP predictor against the number of nodes in the hidden layer based on the macaque dataset.** The legend shows the hidden layer configuration (number of neurons in each hidden layer) as a function of N.

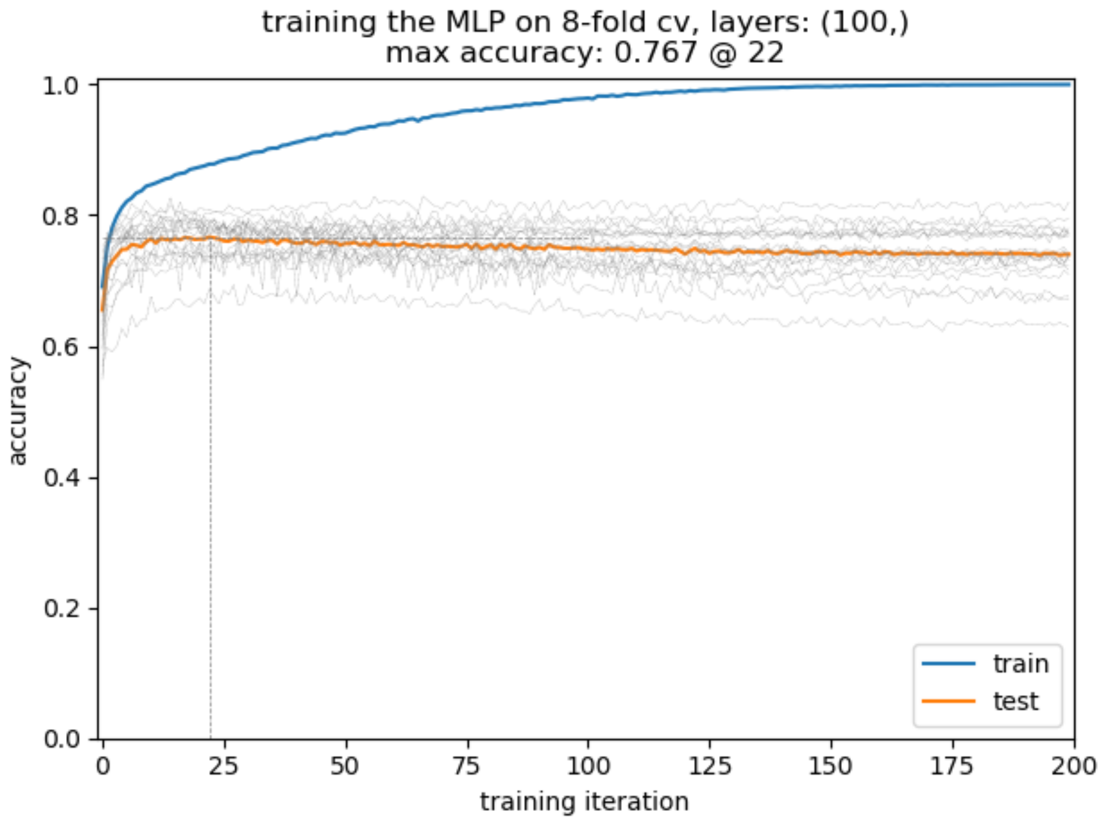


**Fig. S7 Testing overfitting of the MLP predictor against the number of training epochs, macaque dataset.** The gray curves indicate the test set accuracy of individual training runs, the orange curve shows their average. Maximum accuracy on the training set is achieved around 20 iterations.

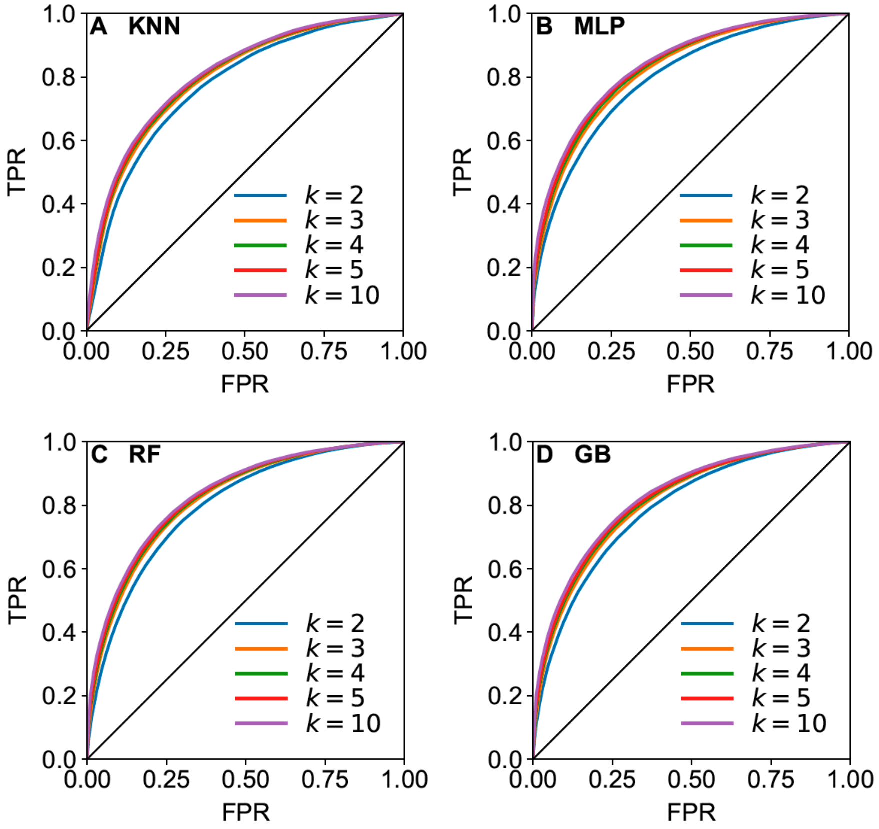


**Fig. S8 ROC curves for binary link prediction using various fold sizes in k-fold cross-validation.** As one can see, $k = 3$ is already sufficient for achieving a close-to-the-best performance. Selecting higher $k$ value, however, decreases the size of the dataset within a fold and increases uncertainty. Throughout we choose to work with $k = 3$. Using the fln-plus-distance feature, averaged over 100 samples.

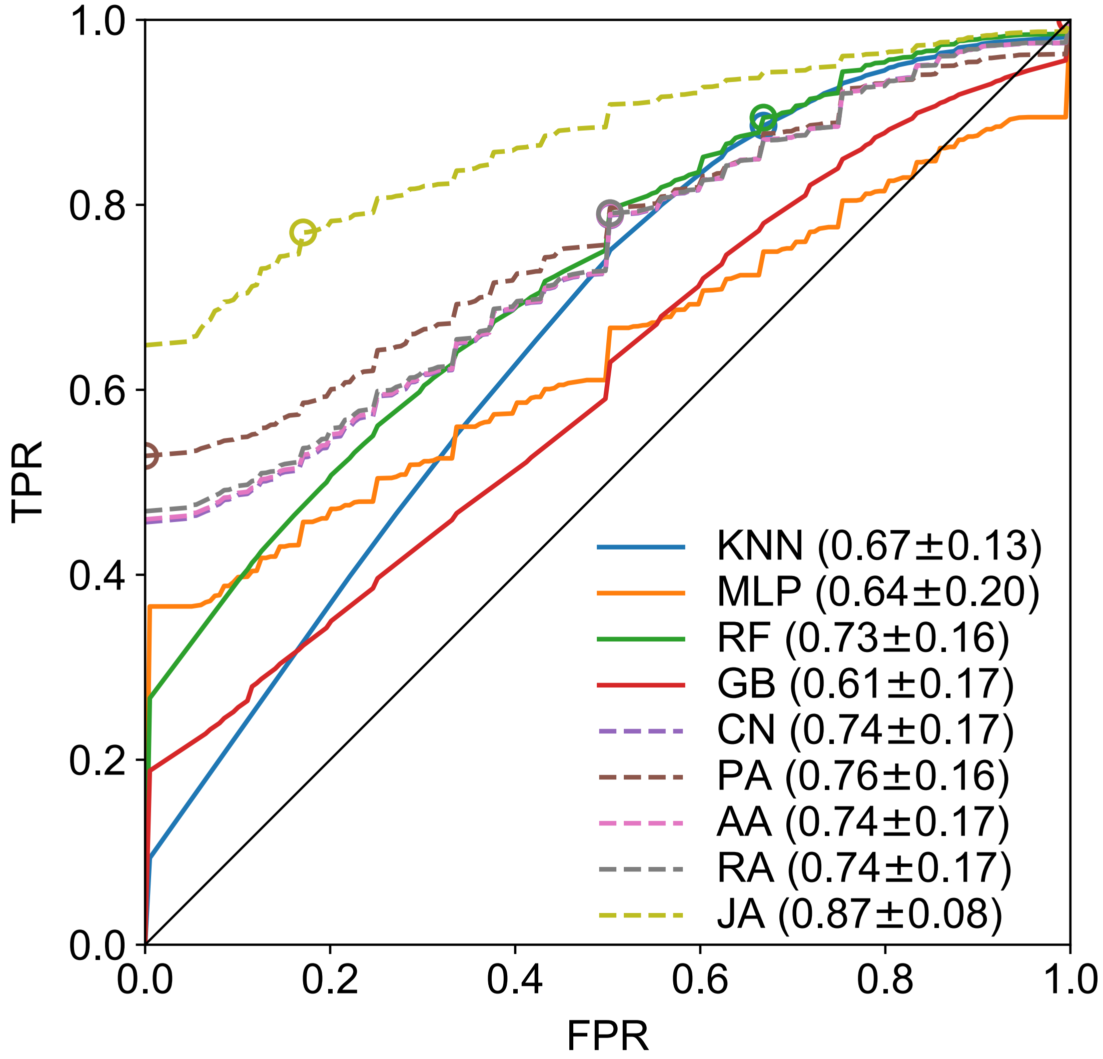


**Fig. S9 ROC curves for binary link prediction in the mouse.** These are based on ML (continuous lines) and CL (dashed lines) algorithms. 3-fold cross-validation, averaged over 100 samples.

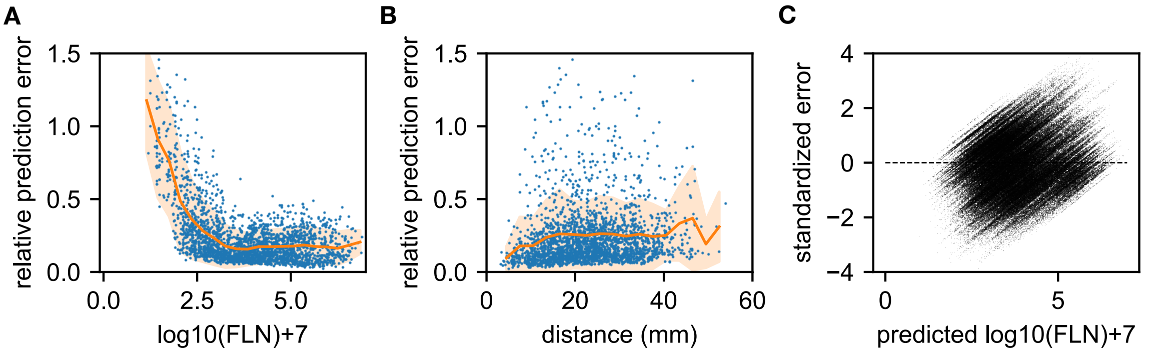


**Fig. S10 Residuals analysis for the Gradient Boosting (GB) algorithm in cross-validation predictions.** Macaque dataset. Based on the fln-plus-distance feature, 3-folded cross-validation, 100 samples. Residual: $Y_{true}(i) - Y_{pred}(i)$, or prediction error. Relative prediction error is $\left|Y_{true}(i) - Y_{pred}(i)\right| / Y_{true}(i)$ where $Y_{true}(i)$ is the true link weight in the data. Predictions are done on the dataset with no-links excluded (so all $Y_{true}(i)$ values are non-zero). Standardized error = $\left(Y_{true}(i) - Y_{pred}(i)\right) / std\_dev$. (A), (B) relative prediction errors (relative absolute residuals) as function of true weights and as function of distance. (C) shows the standardized residuals as function of predicted values. If the points scatter roughly symmetrically around zero along the y-axis, the model is good, most of the pattern/signal has been learned from the data.

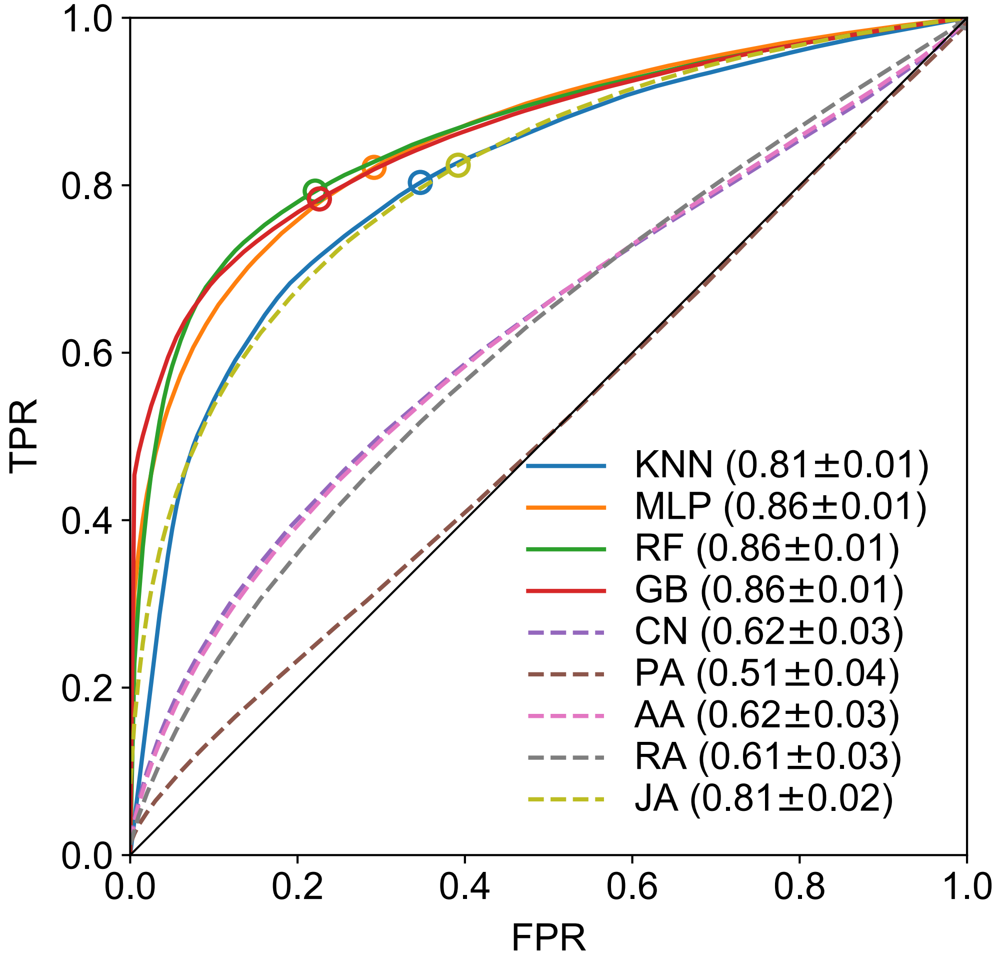


**Fig. S11 ROC curves for binary link prediction in EDR networks**. These are based on the distance matrix of the macaque, and $\lambda = 0.19\ mm^{-1}$ (Ercsey-Ravasz et al., 2013). Using 3-fold cross validation, averaged over 100 samples.

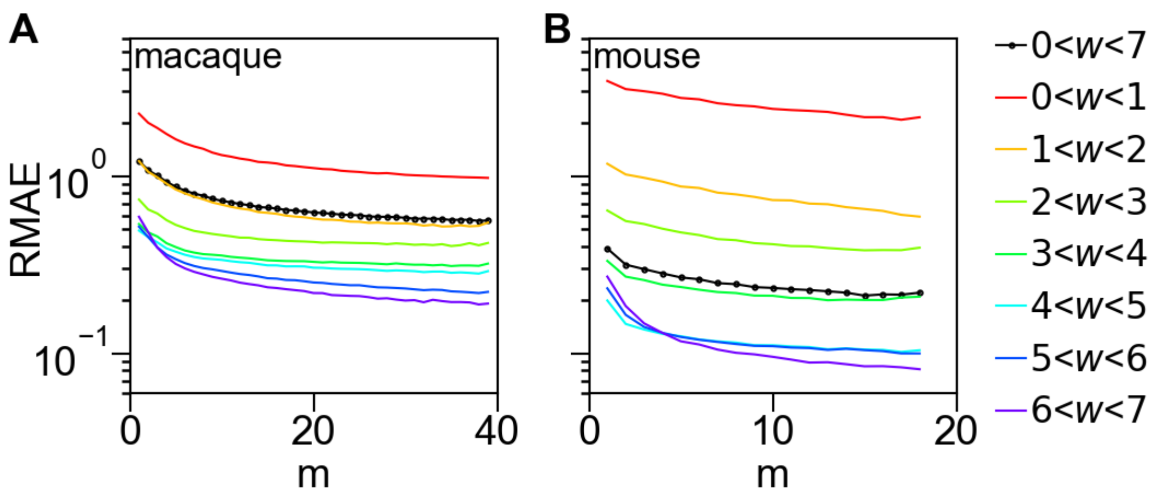


**Fig. S12 Scaling of the external prediction errors.** Here we consider a random subset of $m$ areas from all the targets and instead of leaving an area out of the selected $m$ (as for the case of internal errors shown in Figure 7 of the main text) we train the predictors based on all the data from the $m$ targets to make a prediction for the out-links of *all the others*, *complementary* to $m$ in the total set of targets (i.e., $40 - m$ for the macaque and $19 - m$ for the mouse). These predictions are compared to ground truth and the errors averaged over 500 random target set selections in the same way. We call these the *external relative prediction errors*. Although they look similar to the internal errors shown in Figure 7 of the main text, the values are somewhat different.

### Supplementary Tables

**Table S1. Abbreviations, area names and region assignments for the macaque.**

| Abbreviation | Area name | Region |
|---|---|---|
| 1 | Somatosensory area 1 | Parietal |
| 10 | Area 10 | Prefrontal |
| 11 | Area 11 | Prefrontal |
| 12 | Area 12 | Prefrontal |
| 13 | Area 13 | Prefrontal |
| 14 | Area 14 | Prefrontal |
| 2 | Somatosensory area 2 | Parietal |
| 23 | Area 23 | Cingulate |
| 24a | Area 24, part a | Cingulate |
| 24b | Area 24, part b | Cingulate |
| 24c | Area 24, part c | Cingulate |
| 24d | Area 24, part d | Cingulate |
| 25 | Area 25 | Cingulate |
| 29/30 | Areas 29 and 30 of the retrosplenial cortex | Cingulate |
| 3 | Somatosensory area 3 (includes the primary somatosensory cortex) | Parietal |
| 31 | Area 31 | Cingulate |
| 32 | Area 32 | Cingulate |
| 35/36 | Areas 35 and 36 of the perirhinal cortex | Temporal |
| 44 | Area 44 | Prefrontal |
| 45A | Area 45A | Prefrontal |
| 45B | Area 45B | Prefrontal |
| 46d | Area 46, dorsal part | Prefrontal |
| 46v | Area 46, ventral part | Prefrontal |
| 5 | Somatosensory area 5 | Parietal |
| 7A | Area 7A | Parietal |
| 7B | Area 7B | Parietal |
| 7m | Area 7m | Parietal |
| 7op | Area 7op | Parietal |
| 8B | Area 8B | Prefrontal |
| 8l | Area 8l | Prefrontal |
| 8m | Area 8m | Prefrontal |
| 8r | Area 8r | Prefrontal |
| 9 | Area 9 | Prefrontal |
| 9/46d | Area 9/46d | Prefrontal |
| 9/46v | Area 9/46v | Prefrontal |
| AIP | Anterior intraparietal area | Parietal |
| Core | Auditory core (includes the primary auditory cortex) | Temporal |
| DP | Dorsal prelunate area | Parietal |
| Ento | Entorhinal cortex | Temporal |
| F1 | Frontal area F1 (primary motor cortex) | Frontal |
| F2 | Frontal area F2 | Frontal |
| F3 | Frontal area F3 | Frontal |
| F4 | Frontal area F4 | Frontal |
| F5 | Frontal area F5 | Frontal |
| F6 | Frontal area F6 | Frontal |
| F7 | Frontal area F7 | Frontal |
| FST | Fundus of the superior temporal sulcus | Temporal |

| | | |
|---|---|---|
| Gu | Gustatory cortex | Frontal |
| Ins | Insular cortex | Parietal |
| IPa | Intraparietal sulcus associated area in the superior temporal sulcus | Temporal |
| LB | Belt region of the auditory cortex, lateral part | Temporal |
| LIP | Lateral intraparietal area | Parietal |
| MB | Belt region of the auditory cortex, medial part | Temporal |
| MIP | Medial intraparietal area | Parietal |
| MST | Medial superior temporal area | Temporal |
| MT | Middle temporal area | Temporal |
| OPAl | Orbital periallocortex | Prefrontal |
| OPro | Orbital proisocortex | Prefrontal |
| Pi | Parainsular cortex | Parietal |
| PBc | Parabelt region of the auditory cortex, caudal part | Temporal |
| PBr | Parabelt region of the auditory cortex, rostral part | Temporal |
| PGa | PG associated area of the superior temporal sulcus | Temporal |
| PIP | Posterior intaparietal area | Parietal |
| Pir | Piriform cortex | Temporal |
| ProM | Area ProM (promotor) | Frontal |
| ProSt | Prostriata | Temporal |
| SII | Secondary somatosensory area | Parietal |
| STPc | Superior temporal polysensory area, caudal part | Temporal |
| STPi | Superior temporal polysensory area, intermediate part | Temporal |
| STPr | Superior temporal polysensory area, rostral part | Temporal |
| Sub | Subicular complex | Temporal |
| TEad | Anterior TE, dorsal part | Temporal |
| TEa/m a | Superior temporal sulcus ventral bank area, anterior part | Temporal |
| TEa/m p | Superior temporal sulcus ventral bank area, posterior part | Temporal |
| TEav | Anterior TE, ventral part | Temporal |
| TEO | Temporal area TE, occipital part | Occipital |
| TEOm | Temporal area TE, occipitomedial part | Temporal |
| TEpd | Posterior TE, dorsal part | Temporal |
| TEpv | Posterior TE, ventral part | Temporal |
| TH/TF | Areas TH and TF of the parahippocampal cortex | Temporal |
| Temp.Pole | Temporal pole | Temporal |
| TPt | Temporoparietal area | Temporal |
| V1 | Visual area 1 (primary visual cortex) | Occipital |
| V2 | Visual area 2 | Occipital |
| V3 | Visual area 3 | Occipital |
| V3A | Visual area 3, part A | Occipital |
| V4 | Visual area 4 | Occipital |
| V4t | Visual area 4, transitional part | Temporal |
| V6 | Visual area 6 | Parietal |
| V6A | Visual area 6A | Parietal |
| VIP | Ventral intraparietal sulcal area | Parietal |

**Table S2. Abbreviations, area names and region assignments for the mouse.**

| Abbreviation | Area name | Region |
|---|---|---|
| A | Anterior area | Occipital |
| ACAd | Anterior cingulate area, dorsal part | Cingulate |
| ACAv | Anterior cingulate area, ventral part | Cingulate |
| AId | Agranular insular area, dorsal part | Insular |
| AIp | Agranular insular area, posterior part | Insular |
| AIv | Agranular insular area, ventral part | Insular |
| AL | Anterolateral area | Occipital |
| AM | Anteromedial area | Occipital |
| AUDd | Auditory cortex, dorsal area | Temporal |
| AUDp | Auditory cortex, primary area | Temporal |
| AUDpo | Auditory cortex, posterior area | Temporal |
| AUDv | Auditory cortex, ventral area | Temporal |
| DP | Dorsal posterior area (also known as PD) | Temporal |
| ECT | Ectorhinal area (also referred to as area 36) | Temporal |
| FRP | Frontal pole | Frontal |
| GU | Gustatory area | Insular |
| ILA | Infralimbic area | Frontal |
| LI | Laterointermediate area | Occipital |
| LLA | Laterolateral anterior area | Occipital |
| LM | Lateromedial area | Occipital |
| MM | Mediomedial area | Cingulate |
| MOp | Motor cortex primary | Frontal |
| MOs | Motor cortex secondary | Frontal |
| ORBl | Orbitofrontal area, lateral part | Frontal |
| ORBm | Orbitofrontal area, medial part | Frontal |
| P | Posterior area | Occipital |
| PERI | Perirhinal area (also referred to as area 35) | Temporal |
| PL | Prelimbic area | Frontal |
| PM | Posteromedial area | Occipital |
| POR | Postrhinal area | Occipital |
| PORa | Postrhinal anterior | Occipital |
| RL | Rostrolateral area | Occipital |
| RSPagl | Retrosplenial area, agranular part | Cingulate |
| RSPd | Retrosplenial area, dorsal part | Cingulate |
| RSPv | Retrosplenial area, ventral part | Cingulate |
| SSp-bfd | Somatosensory cortex primary, barrel field | Parietal |
| SSp-lj | Somatosensory cortex primary, lower jaw | Parietal |
| SSp-ll | Somatosensory cortex primary, lower limb | Parietal |
| SSp-nm | Somatosensory cortex primary, nose and mouth | Parietal |
| SSp-tr | Somatosensory cortex primary, trunk | Parietal |
| SSp-ul | Somatosensory cortex primary, upper limb | Parietal |
| SSp-un | Somatosensory cortex primary (unassigned) | Parietal |
| SSs | Somatosensory cortex, secondary | Parietal |
| TEa | Temporal area, anterior part | Temporal |
| TEp | Temporal area, posterior part | Temporal |
| V1 | Primary visual area | Occipital |
| VISC | Visceral area | Insular |

**Table S3. Prediction errors by link weight based only on existing links (w > 0).** Definitions are the same as in Table 1 of the main text. Since the non-links are excluded from the data, the predictors can only predict actual links (cannot predict non-links). Notice, the errors in general are somewhat smaller than in the case when we include the non-links as well.

| | Macaque | | Mouse | | Mac/Mus |
|---|---|---|---|---|---|
| **Non-links excluded** | MAE | RMAE | MAE | RMAE | RMAE ratio |
| Weak ($w_{cut} < w < 3$) | 0.887 | 0.412 | 1.029 | 0.449 | 0.918 |
| Weak-&-Medium ($w_{cut} < w < 5$) | 0.766 | 0.265 | 0.637 | 0.194 | 1.366 |
| Medium-&-Strong ($w > 3$) | 0.786 | 0.176 | 0.555 | 0.124 | 1.419 |
| Strong ($w > 5$) | 0.998 | 0.178 | 0.562 | 0.101 | 1.762 |
| All links ($w > w_{cut}$) | 0.816 | 0.246 | 0.613 | 0.164 | 1.5 |
